# Hippocampal ripples evoke a stereotyped cortical response followed by spindle-mediated network synchronization

**DOI:** 10.64898/2026.05.14.725103

**Authors:** Pin-Chun Chen, Jenny Stritzelberger, Hajo Hamer, Bernhard P. Staresina

## Abstract

Hippocampal sharp-wave ripples (SWRs) are thought to play a key role in systems memory consolidation by broadcasting reactivated memory content to distributed cortical networks while we sleep. Elucidating how this hippocampal-cortical dialogue unfolds at the brain-wide level is therefore essential to understanding how sleep transforms new experiences into lasting memories. Here we combined simultaneous hippocampal intracranial EEG and 21-channel scalp EEG during overnight sleep to characterize the large-scale cortical impact of hippocampal ripple events. We found that individual hippocampal ripples elicited a decodable, phase-locked cortical response at the scalp level. This cortical response was followed by increases in cortico-cortical synchronization and network density in the spindle-band. Mediation analyses revealed a sequential pathway in which ripple magnitude predicted large-scale cortical connectivity through this intermediate cortical response and subsequent spindle activity. These findings demonstrate that hippocampal ripples trigger a two-step cascade, i.e., an early stereotyped cortical response followed by spindle-mediated network synchronization, consistent with the view that SWRs co-activate distributed cortical nodes and potentiate the cortical–cortical connections that support memory consolidation. By demonstrating that ripple-related cortical responses are decodable noninvasively, our results moreover suggest a new strategy for inferring hippocampal ripple activity from scalp electrophysiology.

**Significance Statement:** Hippocampal sharp-wave ripples are brief bursts of coordinated neural activity believed to support memory consolidation by coordinating communication between the hippocampus and the cortex during sleep. Although animal studies show that ripples influence widespread cortical activity, their large-scale cortical effects in humans have previously only been measurable through invasive brain recordings. By combining hippocampal intracranial recordings with scalp EEG, we show that hippocampal ripples evoke a cortical response decodable noninvasively, followed by widespread spindle-mediated synchronization across cortical networks. These findings reveal a temporally structured cascade linking hippocampal activity to large-scale cortical coordination during sleep and suggest that ripple-driven brain-wide responses can be monitored using non-invasive EEG recordings.

## Introduction

Memory consolidation is thought to depend on coordinated interactions between the hippocampus and neocortex, unfolding preferentially during NREM sleep (1). In the systems consolidation framework, newly encoded memories initially rely on the hippocampus and are gradually transferred to distributed cortical sites (2). Thus, reactivated hippocampal traces must be integrated into cortical networks, reshaping connectivity to support long-term storage (3). Arguably the key driver of this hippocampal-cortical dialogue is the hippocampal sharp-wave ripple (SWR) (1). These brief events consist of two distinct components: a large-amplitude slow deflection generated by the synchronous discharge of CA3 pyramidal neurons, upon which fast ripple oscillations (∼80–120 Hz in humans, ∼110–200 Hz in rodents) are superimposed (4). SWRs reflect coordinated activity across hippocampal ensembles and occur predominantly during NREM sleep and quiet wakefulness. In rodents, ripple rates increase following learning and correlate with subsequent memory stabilization (5, 6). Experimental enhancement of ripples strengthens memory consolidation (7, 8), whereas ripple disruption produces learning impairments (9). In humans, intracranial recordings during NREM sleep have linked ripples to declarative memory consolidation (10, 11).

Accumulating evidence across species indicates that SWRs extend their influence far beyond the hippocampus. Ripple-triggered fMRI in anaesthetized and awake macaques shows widespread cortical activation, suggesting rapid reconfiguration of brain states (12, 13). In sleeping rodents, medial prefrontal neurons fire within ∼100 ms of SWRs (14), and Neuropixels recordings in mice reveal ripple-locked changes in firing rates and functional coupling across cortical regions (15). Coordinated hippocampal–prefrontal reactivation is particularly pronounced for larger ripples and can be enhanced experimentally (8). Together, these findings establish that SWRs exert a broad organising influence on cortical activity (16–19). In humans, iEEG studies have begun to characterise hippocampal-cortical interactions during sleep (20–24), but their spatial coverage is constrained by clinical requirements, leaving unclear how SWRs coordinate activity across distributed cortical networks.

The fundamental question, therefore, is what cortical dynamics are set in motion by SWRs, and through which mechanisms are their effects propagated? For the latter, sleep spindles are a strong candidate. Spindles track prior learning-related cortical “hotspots” (25), promote synaptic plasticity by regulating dendritic calcium influx (26), and have been linked to hippocampal-cortical coupling in temporal proximity to SWRs (albeit primarily between the hippocampus and a single scalp site (Cz)) (27). These properties make them well suited to transform transient hippocampal reactivation into longer-lasting cortical changes. Yet, direct characterization of hippocampal-brain-wide interactions has been challenging. Animal studies have mapped ripple-related firing patterns and regional activation, but even dense recordings fall short of full-brain coverage. Moreover, many studies probing large-scale effects have been conducted during wakefulness (15). In humans, knowledge relies largely on iEEG where cortical coverage is sparse and heterogeneous across patients. As a result, we still lack a comprehensive account of how SWRs shape cortical interactions during sleep.

Scalp EEG provides a promising route to address this challenge. While hippocampal ripples themselves are unlikely to be directly observable at the scalp, the cortical responses they evoke should manifest at broader spatial scales (28). Here, we leverage a rare dataset of concurrent hippocampal intracranial EEG and 21-channel scalp EEG recorded during overnight sleep. Hippocampal ripples are identified using stringent criteria and validated via ripple-locked modulation of local neuronal activity. We then test whether individual ripples evoke consistent cortical responses measurable at the scalp, using both univariate and multivariate analyses, and examine how ripples influence cortical connectivity over time. We find that SWRs are followed by a rapid, phase-locked cortical response detectable and decodable in scalp EEG, which is subsequently accompanied by increased spindle-band synchrony and network density peaking ∼200–400 ms post-ripple. Mediation analyses indicate that ripple magnitude predicts later cortical synchronization via this intermediate response and subsequent spindle activity. Together, these findings reveal a two-stage cascade in which ripple-triggered cortical responses precede spindle-mediated network coordination.

## Results

We recorded simultaneous intracranial and scalp EEG from 15 patients with refractory epilepsy undergoing presurgical monitoring (Figure 1A; see Supporting Information S1). Hippocampal SWRs were detected in N2 and N3 overnight sleep (Figure 1B) from macro contacts targeting the hippocampus. Ripple-locked responses were contrasted against surrogate controls (matched non-ripple epochs). Cortical analyses were computed from 21-channel scalp EEG signals time-locked to hippocampal ripple peaks.

**Figure 1.**
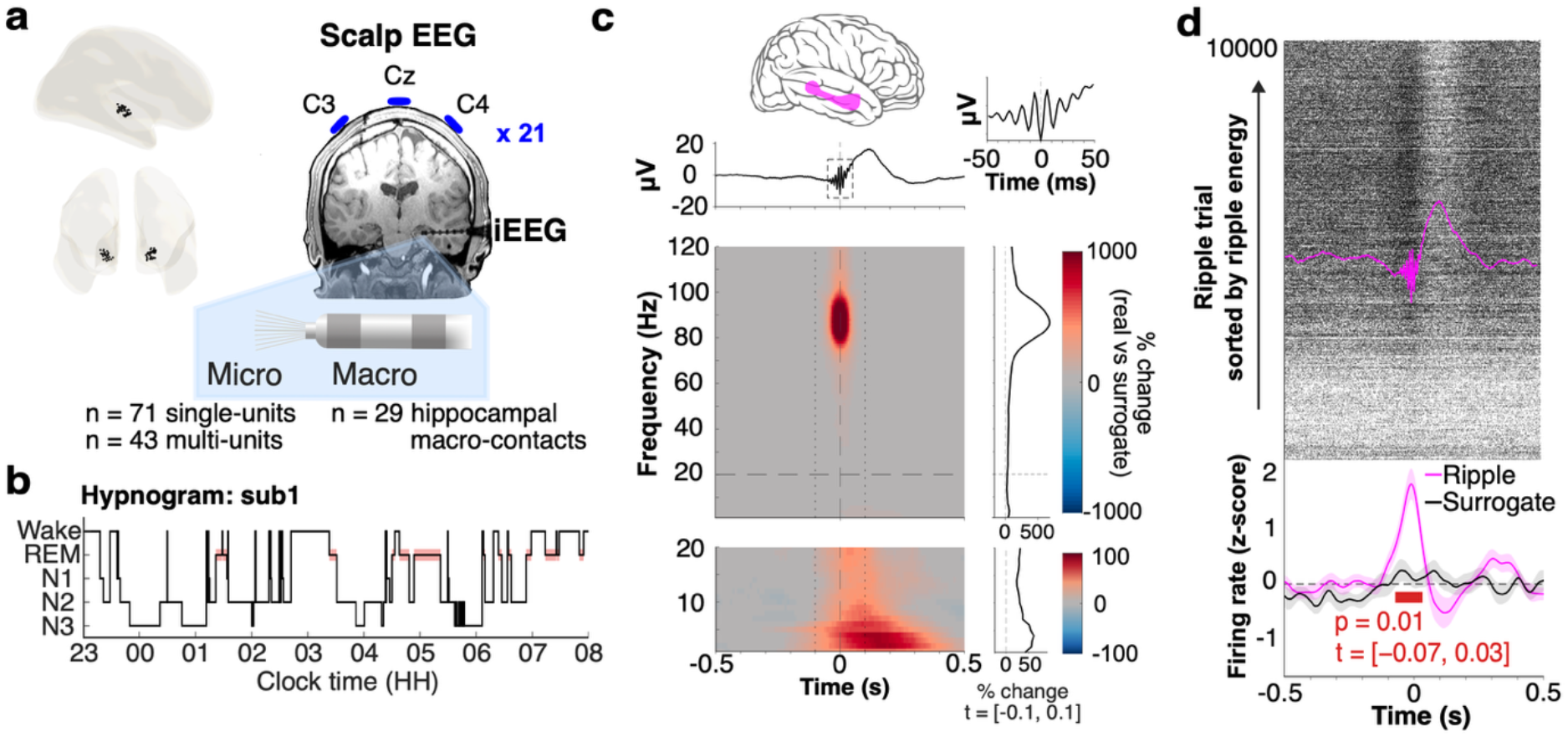
Hippocampal SWRs. (A) Hippocampal recording locations. Left: hippocampal macro contacts from all participants projected onto a standard brain template in MNI space. Right: anatomical location of an example depth electrode targeting the hippocampus, with microwires extending from the electrode tip for single- and multi-unit recordings, and schematic placement of three scalp EEG electrodes (out of a total 21 electrodes placed according to the standard 10-20 montage). **(B) Hypnogram**. Sleep-stage hypnogram from an example participant across the overnight recording. Red segments indicate REM sleep; hippocampal SWRs were detected during NREM sleep stages N2 and N3. **(C) Spectral signature of hippocampal ripples**. Grand-average ripple-locked hippocampal voltage trace (top) and time–frequency representation (bottom). Time–frequency maps were computed for ripples within each hippocampal contact, averaged within contact, and then averaged across contacts (n = 29). The time–frequency decomposition reveals a focal power increase centered at ∼89 Hz, accompanied by a lower-frequency increase in the delta/theta range consistent with the sharp-wave component. **(D) Ripple-locked neuronal firing**. Raster plot showing hippocampal spike activity aligned to ripple peaks across ripple trials (top), sorted by ripple energy (trials with larger energy appear as later trials). The magenta trace indicates the average ripple-locked waveform. Bottom: mean spike firing rate (z-scored across -1 to 1 seconds around ripples) aligned to ripple peaks, pooled across N2 and N3. The red horizontal bar indicates time points of significant firing rate modulation relative to surrogate controls, determined by cluster-based permutation testing (p = 0.010, time = −73 to 32 ms).

Ripple detection was performed on white-matter–referenced macro contact signals using a previously established pipeline (see Methods and Supporting Information S2). Spectral analysis of the hippocampal iEEG signal time-locked to detected ripple events revealed a focal power increase centered at 89 Hz (Figure 1C), accompanied by a prominent low-frequency component consistent with the canonical sharp-wave band of the SWR complex (see Supporting Information S2.4). Ripple energy, quantified as the integral of the ripple envelope between event onset and offset, was used as the primary measure of ripple strength in subsequent analyses.

To further confirm the physiological nature of detected ripple events, we leveraged additional microwire recordings to assess whether hippocampal neuronal firing exhibited the characteristic increase time-locked to ripples (4, 29). Indeed, ripple-locked hippocampal neuronal firing rates increased sharply at ripple onset and peaked near the maximum ripple amplitude before returning to baseline (Figure 1D; cluster corrected permutation test, p = 0.01, time = −73 to 32 ms). These results corroborate that detected events reflect genuine hippocampal SWRs associated with coordinated neuronal population activity.

### Hippocampal SWRs elicit a stereotyped, phase-locked scalp EEG signature

We next examined whether individual hippocampal ripple events produce measurable signatures in simultaneously recorded scalp EEG from 21 electrodes distributed across the scalp (re-referenced to their common average). Event-related potentials (ERPs) time-locked to ripple peaks revealed a multi-component deflection (Figure 2A). Although this response was visually apparent at several channels, cluster-based permutation testing did not identify statistically significant amplitude differences relative to surrogate controls. Similarly, ripple-locked time–frequency representations (TFR; Supporting Information 3.1) of scalp power did not reveal significant power increases or decreases relative to surrogate controls after correction for multiple comparisons (Figure 2B). In contrast to power, however, inter-ripple-trial pairwise phase consistency (PPC; Supporting Information 3.2) revealed two significant clusters of elevated phase locking during ripple relative to surrogate controls (1–5 Hz: p = 0.001; 72– 94 Hz: p = 0.007; Figure 2C), spanning sharp-wave and ripple frequency bands. Thus, ripple-related scalp activity was characterized by consistent phase alignment and more subtle amplitude modulation across trials.

**Figure 2.**
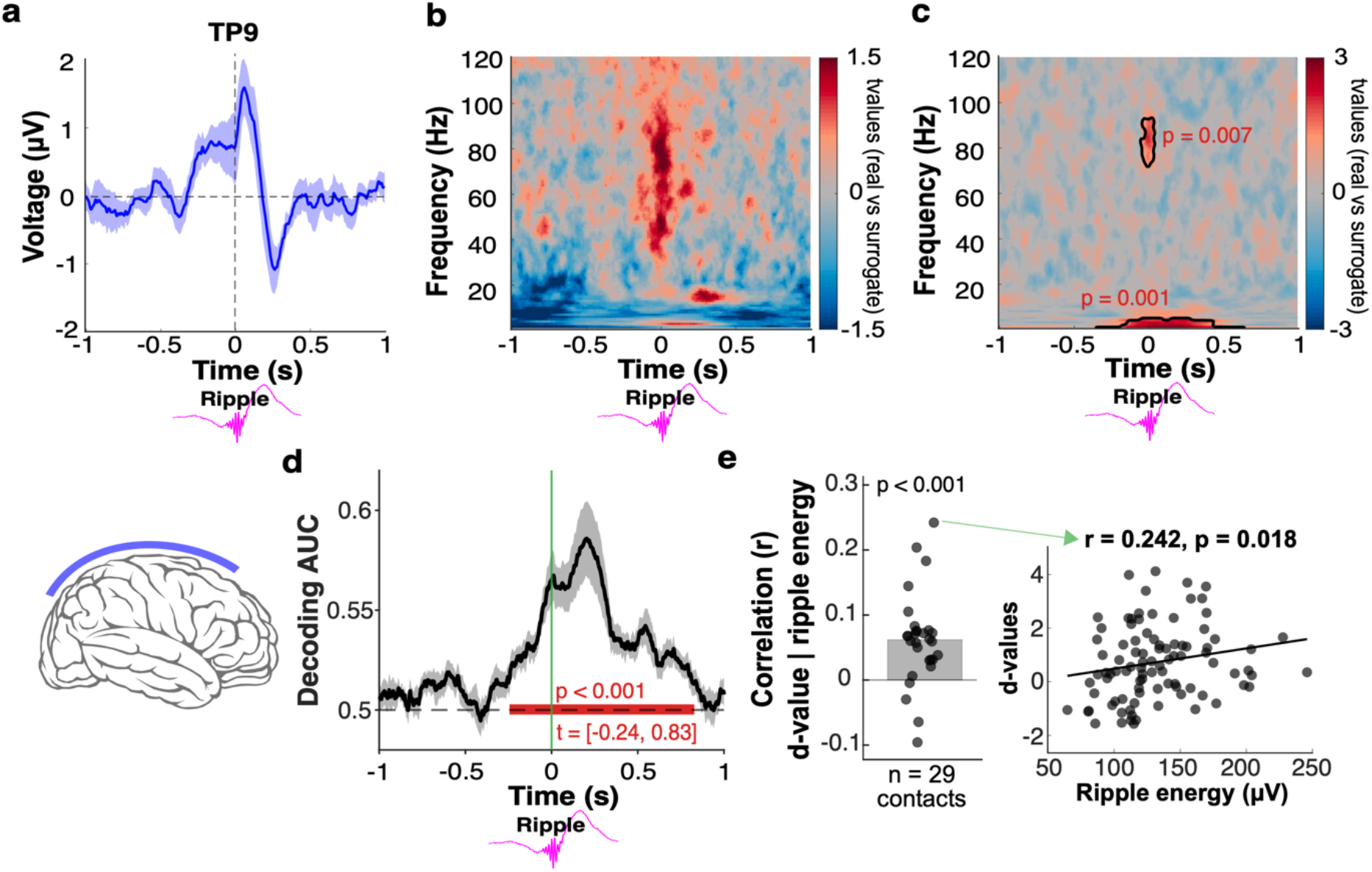
Hippocampal ripples evoke a stereotyped, phase-locked cortical response decodable via scalp EEG. **(A) Ripple-locked scalp event-related potentials (ERP**) at example electrode TP9 (left temporal cortex). Scalp EEG signals were time-locked to hippocampal ripple peaks and baseline-corrected (−1.5 to −1 s relative to ripples). Ripple trials were first averaged within each participant and then averaged across participants to obtain the grand-average ERP (blue trace). Shading indicates the standard error of the mean (SEM) across participants. **(B) Ripple-locked scalp time–frequency power**. Grand-average time–frequency representation (1– 120 Hz) of scalp EEG power aligned to ripple peaks. Power estimates were averaged across scalp electrodes within each participant and averaged at the group level. No significant power differences relative to surrogate controls were detected after cluster-based permutation correction. **(C) Ripple-locked inter-trial phase consistency**. Pairwise phase consistency (PPC) was computed separately for each scalp channel across ripple trials within participants and compared to surrogate controls. Cluster-based permutation testing revealed significant increases in phase alignment in the low-frequency band (1–5 Hz, cluster-corrected *p* = 0.001) and the ripple band (72–94 Hz, cluster-corrected *p* = 0.007). **(D) Decoding ripple-locked scalp responses**. Linear discriminant analysis (LDA) was used to classify ripple-locked epochs versus surrogate controls based on spatial scalp EEG features. Classifiers were trained and evaluated separately for each participant using all ripple trials from that participant, yielding an AUC time course per participant (Figure S1A). The black trace shows the mean classification AUC across participants (± SEM). **(E) Relationship between ripple energy and scalp-based decodability**. Trial-wise LDA decision values were positively associated with ripple energy. Left: each point represents one hippocampal macro-contact; y-axis shows the Spearman correlation between ripple energy and LDA decision value (d-value), with group mean tested against zero by a linear mixed-effects model. Right: example contact showing trial-wise d-values as a function of ripple energy.

To determine whether this scalp EEG response profile allowed above-chance detection of hippocampal ripples at the level of individual events, we applied linear discriminant analysis (LDA) to scalp signals to classify real ripple-locked epochs versus surrogate controls (Supporting Information S3.3). Features were defined as the instantaneous multichannel EEG amplitude vector (21 channels) at each time point. Decoding performance was quantified using the area under the receiver operating characteristic curve (AUC; 0.5 = chance), revealing a significant cluster from −244 to 828 ms relative to ripples (p < 0.001, cluster t-sum = 2420.47), with peak AUC = 0.587 at 200 ms (Figure 2D; Figure S1A), confirming that hippocampal ripples produce reliable, time-locked scalp EEG signatures. When trial averaging was used to boost SNR (n ‘super-trials’ = 10), peak AUC rose to 0.71 (Figure S1C; Supporting Information S3.3.1). Band-specific analyses (Supporting Information S3.3.2) further indicated that decodability was primarily driven by activity in the sharp-wave (2–9.5 Hz) and ripple (80–100 Hz) bands, with no significant decoding observed in the slow oscillation or spindle bands (Figure S2 left). Subtracting the mean evoked response (separately for ripples and surrogate controls) from each trial reduced decoding to chance (Supporting Information S3.3.3; Figure S2 right), indicating that classification depended on a consistent phase-locked response. Importantly, trial-wise LDA decision values (d-values) during peak decoding windows (Figure S1B; Supporting Information S3.3.1) were positively associated with ripple energy in hippocampal macro-contacts (p < 0.001; Figure 2E; Supporting Information S4), demonstrating that larger ripples elicited higher decodability of the cortical response. As an independent validation, ERP-template matching decoding (Supporting Information S3.3.4; Figure S2) also yielded significant above-chance classification, confirming that ripple-locked scalp responses are consistent enough to support template-based classification. Together, these findings establish that hippocampal ripples generate a stereotyped, phase-locked cortical signature whose magnitude tracks ripple strength.

### Hippocampal ripples precede spindle-band cortical synchrony

To examine whether and how SWRs influence cortico-cortical communication, we computed pairwise phase locking values between all 210 possible scalp channel pairs, providing a ripple-trial-level measure of inter-regional phase synchrony (Supporting Information 3.4). For each ripple trial and electrode pair, the phase-locking value (PLV) was computed across 60 time points (500 ms) within each peri-ripple sliding window. Cluster-based permutation testing revealed a significant increase of PLVs for ripple relative to surrogate controls specifically in the spindle (sigma) band (10-17 Hz: p = 0.037; Figure 3A). The effect emerged around the ripple center, peaked between 200 and 500 ms post-ripple, and returned toward baseline by approximately 800 ms. Figure 3B illustrates the time-resolved topography of significant scalp channel pairs. We observed a broad distribution of connections, consistent with widespread cortical synchrony. As a confirmatory control analysis, we computed, using the same approach, the Weighted Phase Lag Index (wPLI (30)), which reduces contributions from zero-lag correlations (likely reflecting volume conduction) and confirmed that real ripples showed significantly greater wPLI than surrogate controls in the significant cluster (mean |wPLI|: 0.486 ± 0.017 vs. 0.477 ± 0.017; paired t-test: t(14) = 6.08, p < 0.001).

**Figure 3.**
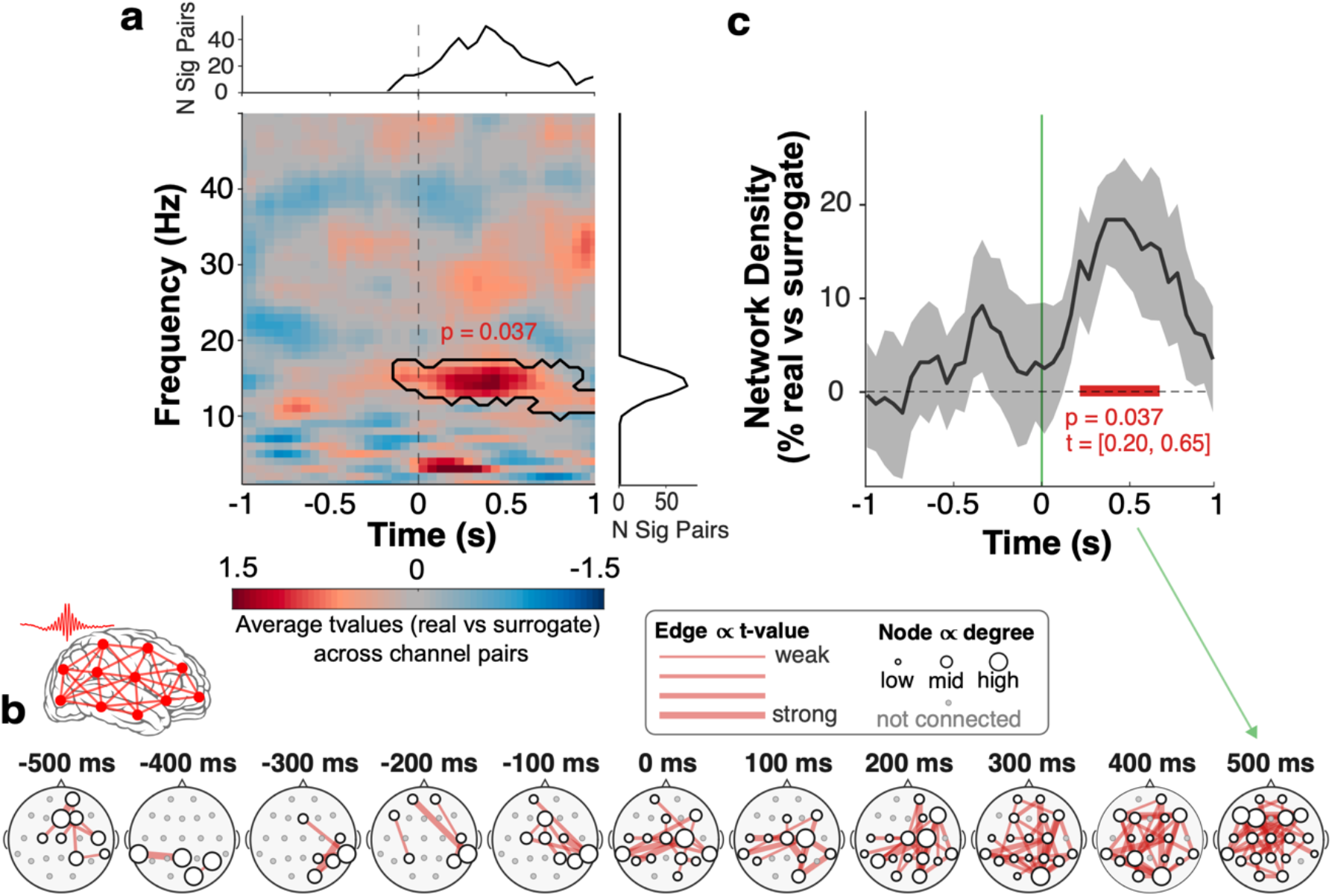
Hippocampal ripples precede spindle-band cortico-cortical synchrony. **(A) Ripple-locked changes in cortical phase synchrony**. Time–frequency representation of pairwise phase-locking values (PLV) averaged across all scalp channel pairs (210 pairs) within participants. PLV was estimated within sliding 500 ms windows proceeding in 50 ms increments across the peri-ripple interval. Cluster-based permutation testing comparing real versus surrogate controls revealed a significant increase in phase synchrony in the spindle band (10–17 Hz, cluster *p* = 0.037). The top panel shows the number of channel pairs contributing to the significant cluster at each time point, and the right panel shows the same quantity marginalised over frequency; both reflect the spatial extent of the effect. **(B) Topography of ripple-associated, synchronized cortical networks**. Scalp connectivity graphs showing spindle-band (12–16 Hz) PLV between electrodes across time windows. Only channel pairs exceeding a t-value threshold of 1.761 (*p* < 0.05, one-tailed, df = 14) are displayed. Edge width scales with the corresponding t-value, and node size reflects channel degree (the number of significant connections that the channel participates in). ∝ … proportional to. **(C) Ripple-associated cortical network density**. Time course of network density derived from spindle-band (12–16 Hz) PLV. Connectivity networks were constructed from 21 × 21 adjacency matrices and thresholded at the top tertile of PLV values. Network density was defined as the proportion of supra-threshold edges relative to all possible connections. Values are shown as change from surrogate controls (mean ± SEM across participants). Red line indicates statistically significant time points (cluster *p* = 0.037).

We next derived graph-theoretic network density from the PLVs (Supporting Information S3.5), applying a threshold retaining the top tertile of the PLV distribution, pooled across all participants, time points, and conditions (ripples and surrogate controls), which yielded fully connected graphs (every node in the network can reach every other node through some path of edges). While Figure 3A displays mean phase coupling strength averaged across all channel pairs, Figure 3B illustrates the spatial layout of significant connections at each time window. Network density complements mean coupling strength by quantifying the proportion of channel pairs exceeding a threshold, an index of how broadly synchrony is distributed across the cortex rather than how strong it is on average. Network density increased significantly from 200 to 650 ms post-ripple (cluster-corrected p = 0.037; Figure 3C), indicating recruitment of a greater proportion of cortical connections relative to surrogate controls. Together, these results demonstrate that hippocampal ripples are followed by a sustained increase of large-scale cortico-cortical communication in the spindle frequency band.

### A two-stage pathway links ripple strength to cortical synchronization

Having identified immediate, transient ripple-locked cortical responses and delayed, sustained spindle-band synchrony, we tested whether these effects form a sequential pathway linking ripple strength to network communication. The temporal dissociation between these responses suggested a possible mechanistic sequence: stronger ripples produce more pronounced cortical signatures, which in turn recruit spindle oscillations that ultimately drive the synchronization of cortical networks.

We formalized this as a three-link cascade mediation model (ripple energy → decoding strength → cortical spindle-band power → cortical spindle-band PLV) using linear mixed-effects models with participant and contact as crossed random intercepts. All three links were significant (ripple energy → decoding: β = 0.067, p < 0.001; decoding → spindle power: β = 0.044, p < 0.001; spindle power → cortical PLV: β = 0.095, p < 0.001), and the indirect effect was significant (a×b×c = 0.0003, 95% CI [0.0001, 0.0005], p = 0.002; Figure 4; Supporting Information S5). Neither the total nor direct effect of ripple energy on cortical PLV was significant (both ps > 0.663), confirming that ripple energy influences cortical synchrony exclusively through the sequential mediating pathway. These findings were confirmed by a structural equation model (lavaan (20)), which yielded consistent path coefficients and a significant indirect effect (a×b×c = 0.0003, 95% CI [0.0002, 0.0005], z = 3.91, p < 0.001; full statistics in Supporting Information S5). Results were further replicated in a control analysis substituting wPLI for PLV (all links and indirect effects significant, ps ≤ 0.002).

**Figure 4.**
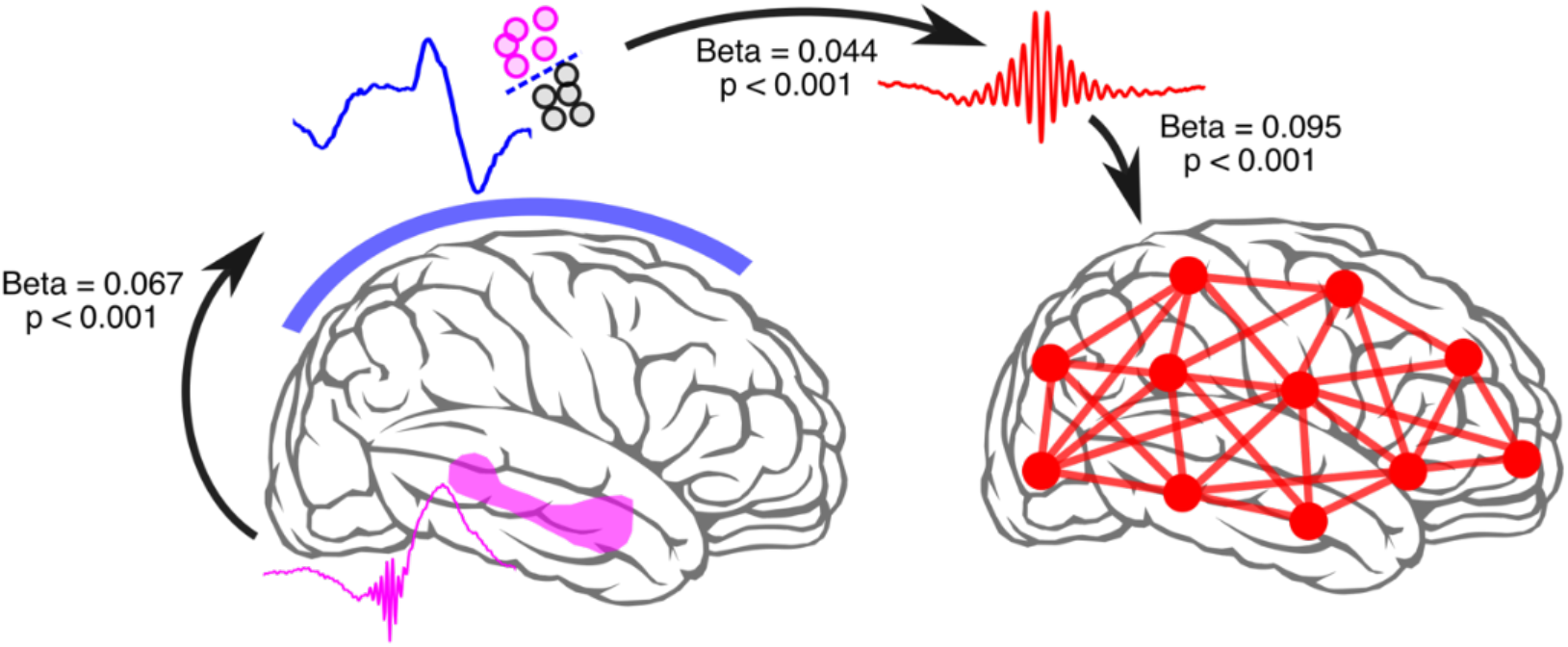
Sequential, two-stage pathway linking hippocampal ripple energy to cortical network synchronization. Mediation analysis testing a ripple-driven cascade linking hippocampal ripples to large-scale cortical communication. Trial-wise ripple energy predicted the decodability of the ripple-locked scalp EEG response, which in turn predicted post-ripple cortical spindle-band power, which predicted cortical spindle-band synchrony (PLV). All three links of the cascade were individually significant, and the sequential indirect effect of ripple strength on cortical synchrony was significant (bootstrap p = 0.002), while the total and direct effects were not, indicating complete mediation through the proposed pathway. Results were replicated using wPLI as a volume-conduction-robust connectivity measure and further confirmed by structural equation modelling. Together, these findings suggest that hippocampal ripples initiate a strength-dependent chain of events, evoking a decodable cortical signature, entraining spindle oscillations, and ultimately synchronizing distributed cortical networks.

Together, these results offer a mechanistic account of how hippocampal ripples influence large-scale cortical dynamics during sleep: individual ripple events trigger a strength-dependent, stereotyped phase-locked cortical response, which in turn entrains spindle oscillations that synchronize cortical networks broadly.

## Discussion

### Ripple-driven two-stage cascade of a hippocampal-cortical dialogue

A central tenet of systems consolidation frameworks is that the function of hippocampal ripples goes beyond coordinating local replay: reactivated hippocampal traces must propagate to distributed cortical sites, enabling the gradual strengthening of cortico–cortical connections to form stable long-term memories (32). Our findings provide systems-level support for this conjecture via combined intracranial and scalp EEG in sleeping humans (Figure 1). Decodability of hippocampal ripple events from scalp EEG revealed a stereotyped cortical response (Figure 2D), which was followed by increased spindle-band cortico-cortical synchrony (Figure 3AB) and network density (Figure 3C). Critically, ripple magnitude predicted both decodability and cortico-cortical synchrony (Figure 4), consistent with the view that hippocampal ripples impact and potentiate distributed cortical nodes, a process that may ultimately forge lasting memory traces.

These observations suggest a two-stage cascade model in which ripple-triggered cortical perturbations reset or reorganize ongoing thalamocortical dynamics. In the first stage, a rapid, phase-locked cortical response emerges across distributed scalp electrodes. This timescale is consistent with rodent studies of hippocampal output, where medial prefrontal neurons fire within ∼100 ms following CA1 activity during SWRs (14), and retrosplenial cortex exhibits similarly rapid ripple-aligned modulation (15, 33). Notably, decoding was not significant in the slow oscillation or spindle bands (see Supporting Information S3.3.2), indicating that the early cortical response does not simply reflect ongoing SO– spindle–ripple coupling. In contrast, band-specific decoding revealed above-chance classifier performance in the sharp-wave range (2–9.5 Hz; Supporting Figure S2B), raising the possibility that the early scalp signal reflects the cortical expression of ripple-associated sharp-wave activity. Consistent with this interpretation, variability in the hippocampal sharp-wave component has been linked to distinct patterns of cortical activation (34), and hippocampal ripples are coupled to cortical sharp waves (35), suggesting that sharp-wave dynamics may shape large-scale cortical responses.

In the second stage, this initial cortical perturbation transitions into spindle-mediated network coordination. It has been shown in rodents that hippocampal SWRs are followed by prefrontal delta waves (0.1-4Hz) that precede spindle activity, producing a ripple-to-spindle delay of ∼500 ms (36). This temporal structure is consistent with a cascade in which hippocampal output first induces a distributed cortical response before recruiting spindle synchronization. At the systems level, this sequence may provide a mechanism by which ripple-reactivated hippocampal content is transformed into widespread cortical plasticity (37). An analogous two-stage process has been observed in humans: Helfrich et al. reported an early ripple-locked prefrontal–MTL interaction followed by a delayed (∼1–2 s) increase in MTL-to-cortex mutual information in the spindle band (20). Our scalp EEG results extend these findings, showing how this cascade manifests as large-scale cortical synchronization measurable noninvasively.

The observed scalp signatures (phase locking and decodable cortical responses) are unlikely to reflect passive volume conduction of hippocampal high-frequency activity. The most direct evidence comes from the mediation analysis: if the scalp signal were simply a propagated copy of the hippocampal signal, it should not account for additional variance in subsequent cortical connectivity effects. Instead, the cortical broadband response and spindle-band activity emerged as necessary intermediates linking ripple energy to large-scale synchronization. Moreover, hippocampal contacts were locally re-referenced to adjacent white matter, while scalp EEG was common-average re-referenced, further reducing the impact of potential signal spread from a hippocampal source.

### Inferring hippocampal ripples from their cortical footprint

A longstanding challenge in human memory research is that hippocampal ripples can only be unambiguously measured invasively, restricting their study to clinical populations. Our findings suggest a potential route to inferring hippocampal ripple activity noninvasively from scalp EEG. While ripple-locked scalp power did not show reliable amplitude modulation (Figure 1B), inter-trial phase consistency revealed significant clustering in both the sharp-wave and ripple frequency bands (Figure 1C). Critically, multivariate decoding of the phase-locked broadband response using a simple linear classifier achieved up to 70% accuracy in distinguishing ripple-from non-ripple epochs (Figure 1D and Supporting Figure S1C). Future work invoking larger iEEG training sets could aim at designing a universal ripple decoder that can be applied to scalp EEG in healthy participants. A complementary strategy may exploit the downstream spindle response. Because cortical spindle-band synchrony peaks at a characteristic delay following hippocampal ripples (Figure 3A), transient increases in spindle-band connectivity could be used as an additional criterion to infer the occurrence of preceding ripple events. Combining these signatures (early phase-locked cortical responses and subsequent spindle-mediated synchronization) may therefore allow the presence of hippocampal ripples to be inferred from scalp EEG, opening new avenues for studying hippocampal memory processing noninvasively in healthy participants.

### Spindles as selective amplifiers of activated cortical circuits

A growing body of work suggests that sleep spindles do not occur randomly or uniformly across the cortex but instead preferentially engage circuits that have previously been biased or activated (38). Recent work has shown that the spatial distribution of spindle activity during post-learning sleep tracks participant-specific cortical activation patterns elicited during learning, indicating that spindles selectively target recently engaged networks (25). Likewise, experimentally biasing cortical excitability prior to sleep using transcranial direct current stimulation shifts the topography of subsequent spindle expression (39), demonstrating that pre-sleep activation states can shape where spindles emerge. Complementary evidence comes from targeted memory reactivation paradigms, where externally presented cues during sleep trigger the reinstatement of specific memory representations, with spindles temporally aligned to periods of increased decodability of the reactivated content (40, 41). Together, these findings converge on the idea that spindles “latch onto” activated cortical circuits, possibly enhancing reactivation and plasticity mechanisms.

Our findings identify hippocampal ripples as a potential additional influence on spindle allocation. Specifically, we show that ripples elicit a stereotyped cortical response which in turn impacts spindle-band synchrony and network density (Figure 3). This finding dovetails with a recent human iEEG study demonstrating that ripple–spindle coupling is linked to memory reactivation (42). In sum, spindles may act on cortical circuits that have been pre-activated—either by prior learning, excitatory brain stimulation, external reminder cues, or, as shown here, by endogenous hippocampal ripple events.

### Limitations and future directions

Although our 21-channel coverage exceeds typical clinical configurations, it remains modest compared with high-density EEG or MEG, hindering the localization of particular cortical hubs that might mediate ripple-locked responses. High-density MEG in healthy participants may further clarify the spatial organization of ripple-driven network dynamics, provided ripples can be reliably detected or inferred (see above). In addition, the present findings are correlational. Future studies combining closed-loop ripple modulation with non-invasive recordings could directly test whether enhancing or suppressing ripples alters large-scale cortical synchronization. Finally, linking ripple-triggered network reconfiguration to behavioral measures of memory consolidation will be essential for corroborating functional relevance.

### Conclusion

The current study shows that human hippocampal SWRs are followed by temporally structured, large-scale cortical dynamics measurable at the scalp. Ripple magnitude predicts subsequent spindle-band synchronization through an intermediate, stereotyped cortical response, supporting a two-stage cascade model of ripple-driven systems coordination. These findings extend animal models of ripple-mediated network communication to non-invasive human electrophysiology and provide a mechanistic bridge between hippocampal output and distributed cortical reorganization during sleep.

## Materials and Methods

### Participants

We analyzed simultaneous intracranial and scalp EEG recordings from 15 patients (7 male; age 19–58 years) with drug-resistant epilepsy undergoing presurgical monitoring at the Epilepsy Center, Erlangen University Hospital, Germany. Each participant provided one overnight recording session (∼8:00 PM– 10:00 AM). Only patients with at least one macro contact located within the hippocampal formation were included. Electrode localization was performed using Brainstorm (43) (see Supporting Information S1.1 and Table S1). The study was approved by the ethics commission of Friedrich-Alexander Universität Erlangen-Nürnberg (142_12B), and written informed consent was obtained in accordance with the Declaration of Helsinki.

### Electrophysiological recordings

Intracranial and scalp EEG signals were acquired simultaneously using an ATLAS recording system (Neuralynx Inc.). Full recording details are provided in Supporting Information S1.

### Intracranial EEG (iEEG)

Recordings were obtained using Behnke–Fried depth electrodes (AdTech Medical, USA) targeting medial temporal structures, with each probe containing macro contacts and a microwire bundle (see S1.1–S1.2). Hippocampal macro contacts were re-referenced to the nearest white-matter contact on the same probe. Signals were acquired at 2048 Hz, bandpass filtered (0.3–200 Hz), notch filtered at line-noise frequencies (50, 100, 150 Hz), and downsampled to 512 Hz for ripple analyses. Across participants, 29 hippocampal macro contacts were included. Microwire spike detection and sorting procedures, including unit inclusion criteria and ripple-locked firing rate analyses, are described in Supporting Information S1.2.

### Scalp EEG

Scalp EEG was recorded from 21 electrodes using the standard 10–20 system with common average re-referencing. Sleep staging was performed manually according to AASM criteria (44). Epochs containing movement artifacts or arousals were excluded. Preprocessing procedures are detailed in Supporting Information S1.3.

### Hippocampal ripple detection

Ripples were detected from white-matter–referenced hippocampal macro-contacts during N2 and N3 sleep using a previously established procedure (27). Prior to detection, signals underwent automated artifact rejection including IED detection (45) (Supporting Information S2.1). Signals were bandpass filtered between 80–120 Hz and the root mean square (RMS) envelope was computed using a 20 ms sliding window. Candidate events exceeding mean + 2.5 SD of the RMS envelope were retained if they met duration (38–200 ms), cycle count (≥3 cycles), and upper amplitude criteria (< mean + 9 SD), and only events exhibiting a clear spectral peak within 80–120 Hz were included (S2.2). Ripple envelope sum (ripple energy) served as the primary measure of ripple strength. Surrogate ripple events were generated using constrained timestamp randomization (S2.3). The dominant low-frequency oscillation co-occurring with ripples was empirically characterized as 2–9.5 Hz via spectral analysis of ripple-locked epochs (S2.4).

### Ripple-locked scalp EEG analyses

Scalp EEG epochs (±2 s) were extracted around hippocampal ripple peaks. Ripple timestamps were pooled across hippocampal contacts within each participant, and matched surrogate controls were generated for comparison. Group-level statistical comparisons between real and surrogate conditions were performed using cluster-based permutation testing, described in the Statistical Analysis section below.

### Time–frequency analysis

Time–frequency representations (TFRs, 1–120 Hz) were computed using Hanning-tapered convolution, retaining complex Fourier coefficients to estimate both power and phase (see Supporting Information S3.1).

### Phase-consistency analyses

Inter-trial phase alignment of ripple-locked scalp activity was quantified using pairwise phase consistency (PPC (46)), a measure of phase clustering across trials that is algebraically unbiased by trial number and therefore permits valid between-participant comparisons when event counts differ (see Supporting Information S3.2).

### Multivariate decoding of ripple-locked scalp activity

To test whether hippocampal ripples produced decodable scalp signatures, linear discriminant analysis (LDA) was applied to classify ripple-locked epochs versus surrogate controls using instantaneous multichannel scalp EEG amplitude (21 channels) as features, using the MVPA-Light toolbox (47). Prior to decoding, EEG signals were smoothed with a 50 ms moving-average kernel. Decoding was performed point-by-point across a ±1 s peri-ripple epoch using 5-fold cross-validation with Ledoit–Wolf shrinkage regularization (48), and performance was quantified as the area under the receiver operating characteristic curve (AUC). Peak decoding latency was identified per participant as the maximum of their smoothed AUC time course within the significant group-level window. Per-trial decision values (d-values), averaged within a ±50 ms window around each participant’s peak latency, were retained for subsequent correlation (S4) and mediation (S5) analyses. A ‘super-trial’ variant, in which trials were averaged in groups of 10 to improve signal-to-noise ratio, was computed as a complementary analysis. Full decoding procedures are described in Supporting Information S3.3.1. They include (i) band-specific decoding across slow oscillation, sharp-wave, spindle, and ripple bands (S3.3.2), (ii) an ERP-removed control in which class-wise evoked responses were subtracted within each cross-validation fold (S3.3.3), and (iii) an ERP-template decoding validation using a sliding-window Pearson correlation classifier (S3.3.4).

### Cross-scalp phase synchrony

Ripple-locked cortical synchrony was quantified using the phase-locking value (PLV) (49) between all scalp electrode pairs, pooled across hippocampal contacts and across N2 and N3 sleep. PLV was computed across time points within a fixed peri-ripple window for each trial, yielding a single per-trial connectivity value per electrode pair, which entered subsequent cascade mediation models as a trial-level outcome variable. PLV was preferred over PPC here because the analysis examined phase consistency across time points within a single trial rather than across trials, i.e. using the same number of phase values throughout to compute the phase locking metric. To address the potential impact of volume conduction, the weighted phase-lag index (wPLI (30)), which is insensitive to zero-phase-lag synchrony, was computed as a confirmatory analysis, averaged within the significant PLV cluster and compared between real and surrogate epochs using a one-tailed paired t-test. Full details are provided in Supporting Information S3.4.

### Cortical network density analysis

To characterize the topology of ripple-associated cortical synchrony, time-resolved functional connectivity networks were constructed from the spindle-band (12–16 Hz) PLV data using a fixed threshold (top tertile of the pooled PLV distribution). Network density was defined as the proportion of surviving edges relative to all possible connections. Full network details are provided in Supporting Information S3.5.

### Ripple attribute correlations

To examine whether ripple strength modulates cortical decodability, per-trial d-values were correlated with ripple amplitude, duration, and envelope energy, using Spearman rank correlations computed within individual hippocampal macro-contacts. Group-level significance was assessed using a linear mixed-effects model with participant as a random intercept, testing the fixed-effect intercept against zero. Full details are provided in Supporting Information S4.

### Cascade mediation analysis

To examine the sequential pathway linking hippocampal ripples to cortical synchronization, a three-link cascade mediation model was implemented: ripple envelope energy → scalp decoding strength (d-value) → scalp spindle-band power → cortical spindle-band PLV. All four variables were estimated on a per-trial basis and z-scored within contact prior to analysis. Spindle-band power was derived from the ripple-locked TFR (S3.1) and PLV from the 500 ms window centred at 300 ms post-ripple, corresponding to the peak of the group-level cluster (S3.4). Each link was estimated as a linear mixed-effects model with participant and contact as crossed random intercepts, following the Baron-Kenny causal steps framework adapted for multilevel data. The indirect effect (product of the three path coefficients) was tested using contact-level bootstrapping (1,000 iterations, 95% CI). Total and direct effects were additionally estimated to confirm full sequential mediation. A control cascade substituting wPLI for PLV as the final outcome was computed to rule out volume conduction. All LME results were confirmed using structural equation modelling with cluster-robust standard errors (31). Full details are provided in Supporting Information S5.

### Statistical analysis

Group-level inference for time–frequency, phase-consistency, decoding, and connectivity analyses used FieldTrip (50) cluster-based permutation testing (1,000 permutations), controlling the family-wise error rate across time, frequency, and channels. All comparisons were performed as dependent-samples t-tests between ripples and surrogate controls. For each participant, ripple trials were pooled across all hippocampal macro contacts prior to computing the effect of interest, such that each participant contributed a single observation per condition. The permutation distribution therefore reflects between-participant variability and accounts for the non-independence of within-participant observations.

Clusters were formed by grouping adjacent data points (participant × channel × frequency × time) exceeding an uncorrected sample-level threshold of p < 0.01. For cross-scalp PLV analyses, channel-pair adjacency was defined by shared electrodes (two pairs were considered adjacent if they shared at least one electrode; S3.4). For network density analyses, clustering was applied across time only. For each cluster, the test statistic was defined as the sum of t-values across all member data points. Cluster-level significance was determined by comparing the observed cluster statistic to the maximum cluster statistic obtained across permutations of condition labels, with results considered significant at a family-wise error rate of α = 0.05. All tests were two-sided, except for decoding AUC (testing above-chance performance) and the wPLI confirmatory analysis (real > surrogate), which were one-sided.

## Acknowledgments

This work is supported by the Marie Skłodowska-Curie Postdoctoral Fellowship (HORIZON-MSCA-2022-PF-01-01; SEP-210878699) awarded to P-C.C. and the European Research Council under the European Union’s Horizon 2020 (grant agreement no. 101001121) awarded to B.P.S.

---

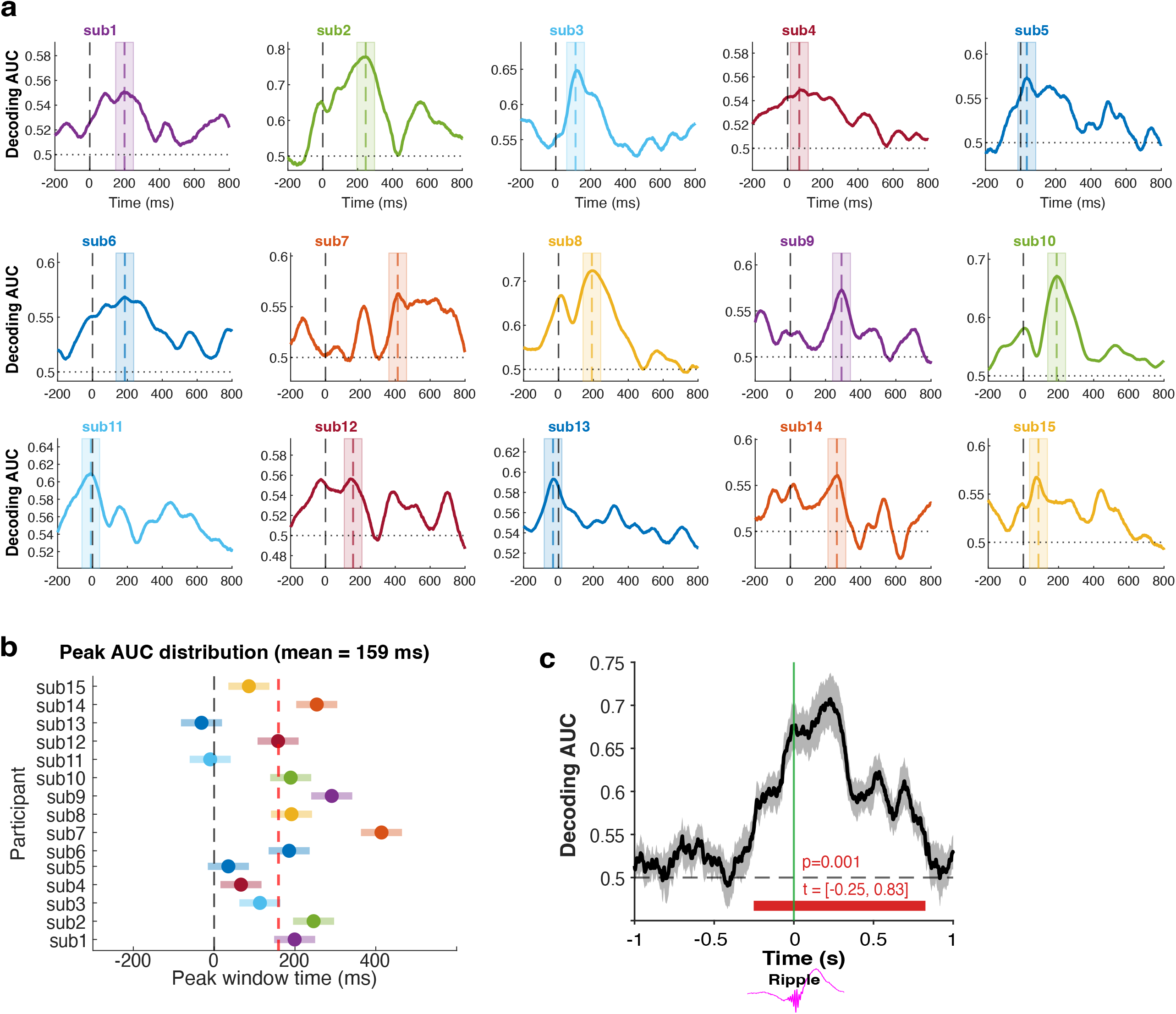

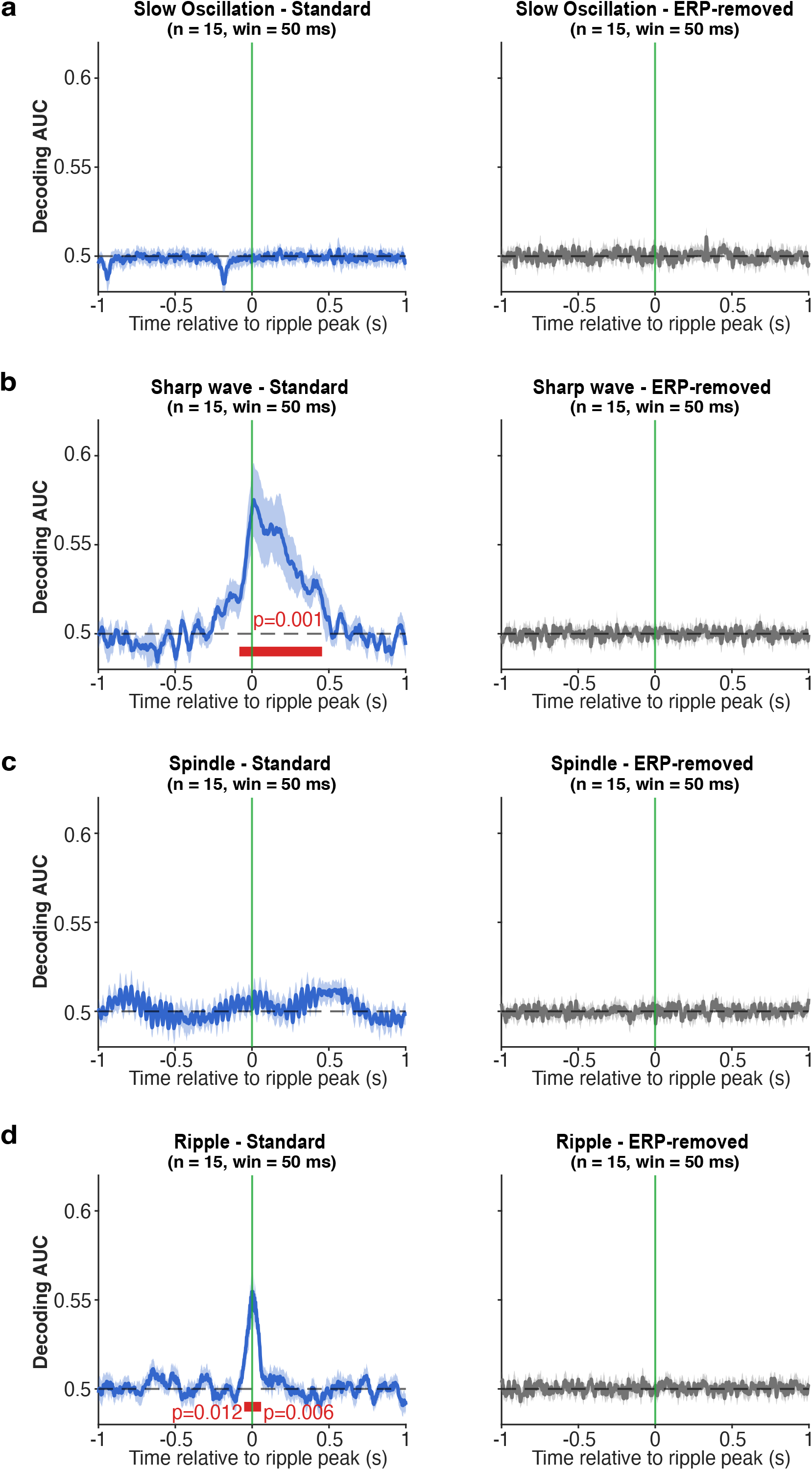

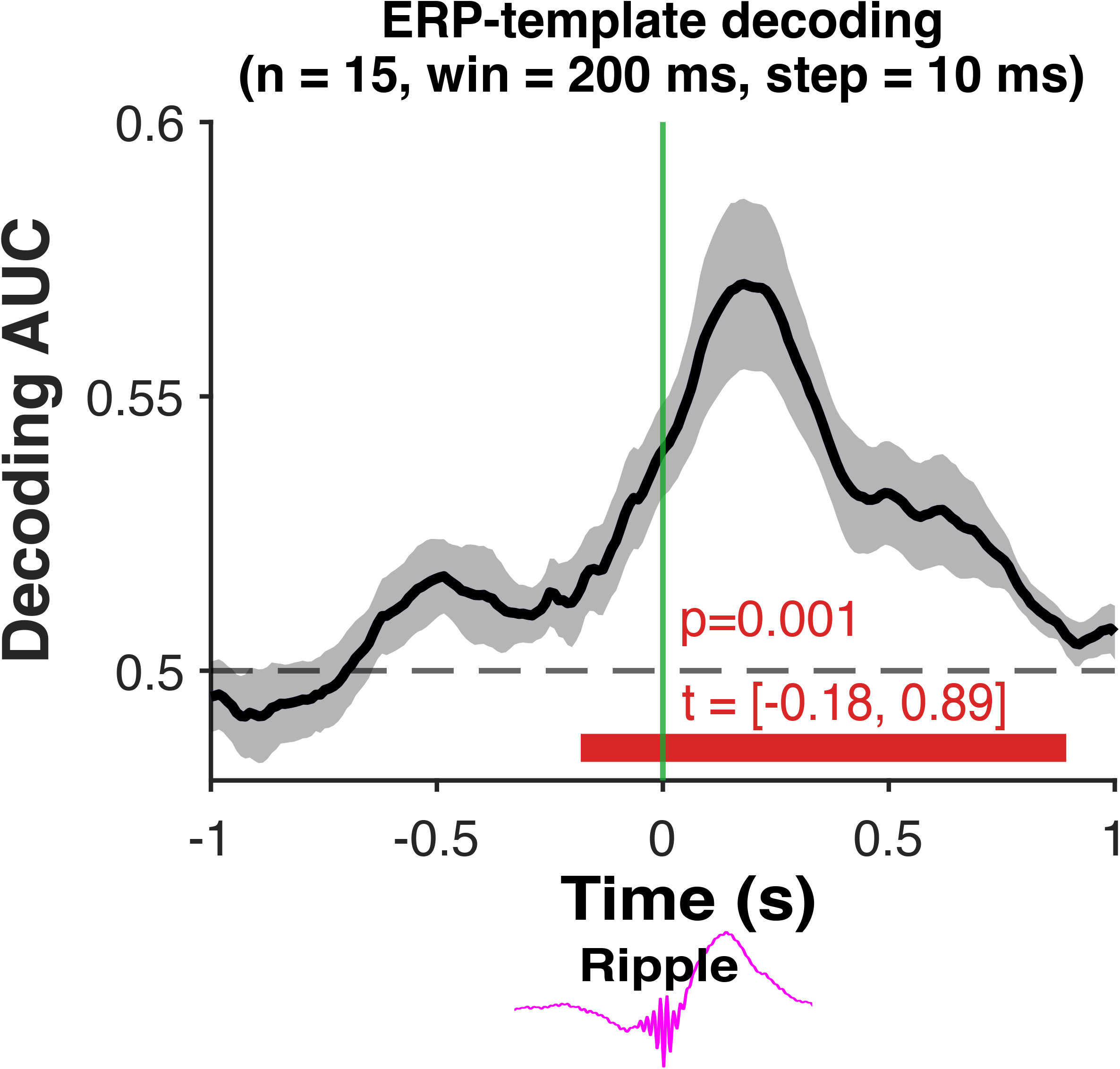

