## Supporting Information for "Hippocampal ripples evoke a stereotyped cortical response followed by spindle-mediated network synchronization"

### Supporting Information for
Hippocampal ripples evoke a stereotyped cortical response followed by spindle-mediated network synchronization.

Pin-Chun Chen,[1,2] Jenny Stritzelberger,[3] Hajo Hamer,[3] and Bernhard P. Staresina[1,4*]

[1]Department of Experimental Psychology, University of Oxford, Oxford, OX1 3EL, United Kingdom

[2]Department of Psychology, Royal Holloway, University of London, Egham TW20 0EX, United Kingdom

[3]Epilepsy Center, Department of Neurology, University Hospital Erlangen, Friedrich-Alexander-University Erlangen-Nürnberg (FAU), 91054, Erlangen, Germany, Full Member of ERN EpiCARE

[4]Centre for Integrative Neuroimaging, Department of Psychiatry, University of Oxford, Oxford, OX3 7JX, United Kingdom

*Correspondence to: Bernhard P. Staresina

**This PDF file includes:**

Supporting text

- S1. Recordings
  - S1.1 Macro-contact recordings
  - S1.2 Microwire recordings and spike sorting
  - S1.3 Scalp EEG recordings
- S2. Hippocampal macro-contact analyses
  - S2.1 Artifact Detection and Rejection
  - S2.2 Ripple Detection
  - S2.3 Surrogate Ripple Events
  - S2.4 Characterization of sharp wave frequency
- S3. Ripple-locked scalp EEG analyses
  - S3.1 Time–frequency representations
  - S3.2 Inter-trial pairwise phase consistency
  - S3.3 Decoding of Ripple-Locked Scalp EEG
    - S3.3.1 Multivariate Pattern Analysis
    - S3.3.2 Band-Specific Decoding
    - S3.3.3 ERP-Removed Decoding Control
    - S3.3.4 ERP-Template Decoding Validation
  - S3.4 Cross-scalp phase synchrony
  - S3.5 Network density analysis
- S4. Ripple Strength vs Cortical Decodability
- S5. Cascade Mediation Analysis

Figures S1 to S3
Tables S1 to S3
SI References

**Supporting Information Text**

#### S1. EEG Recordings

Intracranial recordings were obtained using Behnke–Fried depth electrodes (AdTech Medical, USA) targeting medial temporal lobe structures. Signals were acquired using an ATLAS recording system (Neuralynx Inc.). Each probe contained macro-contacts (1.3 × 1.6 mm) and a microwire bundle consisting of eight microwires (40 μm diameter) and one reference wire.

##### S1.1 Macro-contact recordings

Macro-contact signals were referenced to the nearest white-matter contact on the same probe to reduce volume conduction and shared noise. Across participants, 29 hippocampal macro-contacts were identified within the hippocampal formation and included in the analyses. Macro-contact signals were sampled at 2048 Hz. Data were subsequently bandpass filtered between 0.3 and 200 Hz, downsampled to 512 Hz, and notch filtered at 50 Hz and its harmonics (100 and 150 Hz) to remove line noise. Within each participant, contacts located within the hippocampal formation were identified via visual inspection of postoperative T1-weighted anatomical MRI scans. These macro-contact channels were used for ripple detection and all hippocampal iEEG analyses. For visualization, electrode coordinates were transformed to MNI space using the Brainstorm toolbox (1) (Fig. 1A; Table S1).

##### S1.2 Microwire recordings and spike sorting

Microwire signals were sampled at 32,768 Hz and used for spike detection and sorting to validate ripple events. Spike extraction and clustering were performed using Combinato (2). Negative voltage deflections were extracted as candidate spikes. Automated artifact removal was applied using the DER algorithm (3). Cluster assignments were manually validated by an experienced rater, and artifactual clusters were removed. Because automated clustering in Combinato can over-segment spike data, clusters were manually merged based on waveform similarity, cross-correlograms, and firing characteristics. Units with a mean firing rate below 0.05 Hz across the N2 and N3 period were excluded. In total, 71 single units and 43 multiunits were identified across hippocampal microwires. For ripple-locked firing rate analyses, firing rates were calculated by pooling all units (single and multi-units) at each macro-contact level. Statistical comparisons between ripple-locked and surrogate firing rates were performed using a cluster-based permutation test at the macro-contact level (Figure 1C).

##### S1.3 Scalp EEG recordings

Scalp EEG was sampled at 2048 Hz, recorded from 21 electrodes placed according to the international 10–20 system (Fp1, Fp2, F7, F3, Fz, F4, F8, T7, C3, Cz, C4, T8, P7, P3, Pz, P4, P8, O1, O2, TP9, TP10). One participant lacked TP9 and TP10 recordings; these channels were padded with NaN values for that participant. Signals were re-referenced using a common average reference (CAR) computed across all 21 scalp channels to reduce shared noise and volume-conducted effects. For sleep staging, scalp EEG was downsampled to 128 Hz and bandpass filtered between 0.3 and 50 Hz. Sleep stages were scored manually according to AASM criteria. Epochs containing movement artifacts or arousals were excluded from all analyses. For ripple-locked analyses, scalp EEG data were downsampled to 512 Hz to match the sampling rate of the hippocampal recordings.

#### S2. Hippocampal macro-contact analyses

##### S2.1 Artifact Detection and Rejection

To minimize contamination from interictal epileptiform discharges (IEDs) and other artifacts, automated artifact detection was applied independently to each white-matter-referenced hippocampal macro-contact prior to ripple detection, followed by visual confirmation. Signals were z-scored separately within each sleep stage to account for differences in physiological signal properties across vigilance states. For each channel and sleep stage, time points were marked as artifactual if any of the following measures exceeded a z-score threshold of 4:

- absolute amplitude of the raw signal
- absolute amplitude of the first temporal derivative of the signal (capturing sharp transients characteristic of IEDs)

IEDs were additionally detected using an automated algorithm (4). Flatline segments, defined as five or more consecutive samples with zero amplitude, were also marked as artifactual. Detected artifact segments were padded by ±0.5 s to conservatively exclude pre-, peri- and post-artifact contamination. Artifact-free segments shorter than 1s were subsequently labelled as artifactual to avoid fragmentation of clean data intervals. Artifact-free sleep durations of each white-matter-referenced hippocampal macro-contact in each sleep stage are reported in Table S2.

##### S2.2 Ripple Detection

Ripples were detected independently for each hippocampal macro-contact channel and separately for N2 and N3 following an established pipeline (5). Only artifact-free data were included. Signals were bandpass filtered between 80 and 120 Hz using a zero-phase finite impulse response (FIR) filter. The root mean square (RMS) envelope of the filtered signal was computed using a 20 ms sliding window and smoothed with a 20 ms moving average. For each channel and sleep stage, ripple candidates were defined as contiguous periods in which the RMS envelope exceeded a threshold of mean +2.5 standard deviations calculated from artifact-free samples within that stage. Events exceeding mean +9 standard deviations were excluded as likely artifacts. Candidate events shorter than 38 ms or longer than 200 ms were rejected. Events containing fewer than three oscillatory cycles were also excluded.

To further reduce false positives, each candidate ripple underwent spectral validation. A time–frequency representation was computed within a ±250 ms window (65–135 Hz, 2 Hz resolution) using adaptive Hanning-tapered windows. Power was averaged within ±50 ms around the RMS peak and normalized by its maximum value. Spectral peaks were identified using MATLAB's *findpeaks* function with a minimum prominence of 0.2. Events were retained only if at least one spectral peak fell within the ripple band (80–120 Hz). Ripple attributes are reported in Table S3 for descriptive characterization.

##### S2.3 Surrogate Ripple Events

To provide a statistical baseline for ripple-locked analyses, surrogate control events were generated for each participant by randomly shifting detected ripple timestamps within a 300 s search window centered on each real ripple, constrained to artifact-free samples within the same sleep stage. For each real ripple, five candidate surrogate timestamps were generated, and any candidate falling within 1 s of a real ripple event was excluded to prevent contamination of the surrogate distribution with genuine ripple-related activity. The final number of surrogate trials was matched to the number of real ripple trials by random subsampling. This procedure yields surrogate events that are matched to real ripples in terms of sleep stage and approximate time of night. Importantly, no restriction was placed on the occurrence of sleep spindles within surrogate epochs, ensuring that any spindle-band effects observed in real relative to surrogate ripple epochs reflect a genuine temporal association with ripple events rather than a general suppression of spindle activity in the surrogate condition.

##### S2.4 Characterization of Sharp-wave Frequency

To empirically characterize the dominant low-frequency oscillation during ripple events, power spectral density was computed from ±500 ms windows centered on each detected ripple for both real and surrogate epochs. Trial-averaged spectra were estimated using a multitaper fast Fourier transform with a Hanning taper (frequency resolution: 0.5 Hz; range: 2–30 Hz). Two complementary approaches were applied to isolate frequency-specific power increases in real relative to surrogate epochs. In the first, the aperiodic (1/f) component was removed using the specparam algorithm (6), fitted to the average of the real and surrogate spectra to obtain an unbiased slope estimate; both spectra were then flattened against the same aperiodic fit. In the second, the log ratio of real to surrogate power ($\log_{10}$[P_real / P_surrogate]) was computed per contact, directly canceling the common aperiodic component and any broadband amplitude differences without requiring explicit model fitting. For each approach, the resulting spectra were entered into a cluster-based permutation test across all hippocampal contacts (1,000 permutations; dependent-samples t-test; two-tailed; cluster $\alpha = 0.05$). Both approaches consistently identified a significant positive cluster spanning 2–9.5 Hz, establishing this range as the empirical sharp-wave frequency band used in subsequent decoding analysis in S3.3.2.

#### S3. Ripple-Locked Scalp EEG Analyses

##### S3.1 Time-Frequency Analysis

Time–frequency representations (TFRs) were computed for each scalp channel using Hanning-tapered convolution implemented in FieldTrip. Analyses covered 1–120 Hz with estimates calculated every 10 ms within a ±1s window centered on ripple events. Frequency-dependent window lengths were used: for frequencies ≤5 Hz, the number of cycles equaled the frequency value; for higher frequencies, windows contained at least five cycles with a minimum duration of 100 ms. Complex Fourier coefficients were retained, enabling both power and phase analyses. Group-level differences between real and surrogate ripple TFR power were evaluated using the cluster-based permutation framework described in the Statistical Analysis section.

##### S3.2 Inter-Trial Pairwise Phase Consistency

To assess ripple-locked phase alignment across trials, pairwise phase consistency (PPC) was computed for each scalp channel using the instantaneous phase of the complex Fourier spectra (7). PPC provides a bias-free measure of inter-trial phase clustering that is independent of trial count.

$$PPC = \frac{|\sum_{k=1}^{N} e^{i\phi_k}|^2 - N}{N(N-1)}$$

where $\phi_k$ is the instantaneous phase of trial $k$ and $N$ is the number of trials. The same procedure was applied to surrogate ripple epochs to provide a baseline estimate of phase consistency expected by chance. Group-level differences in PPC between real and surrogate ripple epochs were assessed using the cluster-based permutation framework described in the Statistical Analysis section.

##### S3.3 Decoding of Ripple-Locked Scalp EEG

###### S3.3.1 Multivariate Pattern Analysis

To assess whether hippocampal ripples produce detectable signatures in simultaneously recorded scalp EEG, we employed a multivariate pattern analysis (MVPA) decoding approach. For each participant, scalp EEG epochs time-locked to detected ripples were contrasted against epochs time-locked to surrogate controls using a regularized linear discriminant analysis (LDA) classifier, implemented in the MVPA-Light toolbox (8). Where multiple hippocampal contacts were available, epochs from all contacts were pooled prior to decoding. Trials were balanced between ripples and surrogate controls, capped at 2,000 trials each. Prior to decoding, EEG signals were smoothed using a 50 ms moving-average kernel. Decoding was performed in a point-by-point fashion across the ±1 s peri-ripple epoch, employing the '*mv_classify_across_time'* function in the MVPA-Light toolbox. At each time point, the instantaneous multichannel EEG amplitude vector (21 channels) was used as the feature vector.

Classification used 5-fold cross-validation with LDA regularized via Ledoit–Wolf analytical shrinkage (using MVPA-Light's *reg='shrink', lambda='auto'* setting (7)), which automatically estimates an optimal regularization parameter to stabilize the covariance matrix. At each fold, features were z-scored using the training-fold mean and standard deviation before classification. The discriminant axis estimated from training data was applied to held-out trials, projecting each trial's feature vector to a scalar decision value (*d-value*): positive values indicate classification as real ripples, negative values as surrogate controls. Decoding performance was quantified as the area under the receiver operating characteristic curve (AUC), computed from cross-validated decision values using the Mann–Whitney U statistic.

Per-trial d-values from all five held-out sets were averaged and linked to trial-level ripple attributes for subsequent correlation (S4) and mediation (S5) analyses. Specifically, to obtain a single scalar decoding index per trial, d-values were averaged within a ±50 ms window centered on each participant's peak decoding latency. Peak latency was identified per participant as the maximum of their smoothed AUC time course (50 ms moving average) within the group-level above-chance decoding window (−244 to 828 ms relative to ripple peak; Figure 2D; individual-participant decoding shown in Figure S1A). D-values were then averaged within a ±50 ms window centered on each participant's peak latency (Figure S1B), yielding a single scalar decoding strength per trial for subsequent correlation (S4) and mediation (S5) analyses.

To complement the main single-trial result, we repeated the decoding using 'super-trial' averaging to improve SNR. Within each hippocampal contact, trials were randomly assigned to non-overlapping groups of 10 and averaged to yield a single super-trial per group, separately for real-ripple and surrogate-control epochs to prevent mixing of class labels. 'Super-trials' were then pooled across contacts and passed to the same LDA pipeline described above. The 'super-trial' analysis yielded a markedly improved peak AUC of 0.71 at 228 ms, with a significant above-chance cluster from −252 to 826 ms (cluster p = 0.001, t-sum = 2810.37; Figure S1C), closely mirroring the temporal profile of the main single-trial result in Figure 2D.

###### S3.3.2 Band-Specific Decoding

To examine the spectral contributions to ripple-locked scalp signals, decoding was repeated on signals filtered within four canonical frequency bands prior to feature extraction: slow oscillation (0.5–1 Hz; Figure S2A, left), sharp-wave (2–9.5 Hz; Figure S2B, left), spindle (12–16 Hz; Figure S2C, left), and ripple band (80–100 Hz; Figure S2D, left). Bandpass filtering used a 4th-order Butterworth filter applied with zero-phase forward-backward filtering (MATLAB filtfilt). Following filtering, each band's signal was smoothed with a 50 ms moving-average kernel before decoding, which followed the same pipeline described in S3.3.1.

Significant above-chance decoding was observed in the sharp-wave (2–9.5 Hz), and ripple (80–100 Hz) bands, but not in the slow oscillation (0.5–1 Hz) or spindle (12–16 Hz) bands (Figure S2, left panels). In the sharp-wave band, significant decoding was observed from −82 to 456 ms (cluster p = 0.001, t-sum = 846.77), with a peak AUC of 0.575 at 12 ms. In the ripple band, significant decoding was confined to brief peri-event windows, comprising two clusters: one from −52 to 2 ms (cluster p = 0.012, t-sum = 87.98, peak = 0.555 at −2 ms) and another from 6 to 58 ms (cluster p = 0.006, t-sum = 111.46, peak = 0.553 at 10 ms).

**S3.3.3 ERP-Removed Decoding Control**
To determine whether decoding performance reflected phase-locked evoked responses, an ERP-removed control variant was computed for each band. Within each training fold, the class-wise mean feature vector was subtracted from every trial of the corresponding class before classifier training; the same fold-derived class means were then subtracted from test-fold trials before prediction. This strictly-within-fold procedure ensures that the classifier has no access to the average evoked response and can only exploit residual trial-to-trial variability, with no leakage of test-set information into the subtraction. Following ERP removal, decoding AUC collapsed to chance across all frequency bands (Figure S2, right panels). This pattern indicates that the scalp EEG signature of hippocampal ripples is carried predominantly by a consistent phase-locked evoked response.

**S3.3.4 ERP-Template Decoding Validation**
As an independent validation of ripple-locked scalp EEG patterns, we implemented a template-matching decoding approach. Scalp EEG epochs time-locked to real and surrogate ripples were pooled across hippocampal contacts and balanced to equal class sizes following the same procedure as the LDA analysis (S3.3.1).

Classification was performed using 5-fold cross-validation. Within each training fold, class-specific ERP templates were computed by averaging all training trials belonging to each class separately, yielding one mean spatiotemporal pattern per class (channels × time). For each test trial, classification was based on Pearson correlation between the test trial's spatiotemporal pattern and each class template (the trial was assigned to the class whose template yielded the higher correlation). Decoding was performed in a sliding-window fashion across the ±1s peri-ripple epoch using a 200 ms window in 10 ms steps.

Group-level cluster-based permutation test revealed a significant decoding from -182 to 893 ms relative to ripple peak (p = 0.001; peak AUC = 0.571 at 180 ms; Figure S2), confirming that the scalp EEG signature of hippocampal ripples is sufficiently consistent in its spatiotemporal morphology to support classification by template similarity alone. The convergence of this result with the LDA decoding, obtained through a fundamentally different classification approach, strengthens the conclusion that ripples are associated with a reliable, phase-locked cortical evoked response.

**S3.4 Cross-Scalp Phase Synchrony**
Ripple-associated changes in cortical synchrony were quantified using phase-locking values (PLV) between all scalp electrode pairs. Complex Fourier spectra were obtained as described in S3.1, restricting analyses to 1–50 Hz. For each channel pair and trial, instantaneous phase differences were computed at each frequency and time point from the complex Fourier coefficients, and PLV was calculated as the mean resultant length of the phase difference vector (*circ_r* from the circstat toolbox (10)). Phase locking values were averaged within a sliding 500 ms window stepped in 50 ms increments across ±1.25 s relative to ripple center, with each window comprising 60 timepoints, over which phase consistency was estimated. Per-participant estimates were obtained by averaging across all valid ripple trials, pooling across hippocampal contacts and across N2 and N3. Group-level differences between real and surrogate PLV were assessed using the cluster-based permutation framework described in Statistical Analysis, with channel-pair adjacency defined by shared channels (two channel pairs were considered adjacent if they shared at least one electrode, e.g. Fz–Cz and Fz–Pz are adjacent via Fz). As an additional control for volume conduction, cortical phase coupling was quantified using the weighted phase-lag index (wPLI). Because the imaginary component of the cross-spectrum is zero for

signals with no phase lag, wPLI is insensitive to zero-phase-lag synchrony likely arising from volume conduction. Specifically, wPLI weights each observation by the magnitude of the imaginary component of the cross-spectrum, such that phase relationships near 0° or 180° contribute less to the estimate. wPLI was computed across time points within each trial rather than across trials, yielding a single per-trial connectivity value. For the confirmatory test, wPLI was averaged within the significant PLV cluster (across all channel pairs, frequencies, and time windows belonging to the cluster) separately for real and surrogate epochs, and the resulting per-participant scalar values were compared using a one-tailed paired t-test (real > surrogate).

For visualization, t-values were averaged across channel pairs to yield a reduced time–frequency summary map, with significant clusters overlaid as contours. Spindle-band (12–16 Hz) connectivity was additionally displayed as scalp graphs per time window, retaining only pairs exceeding t > 1.761 ($p < 0.05$, one-tailed, df = 14). Edge width scales with t-value and node size reflects channel degree (the number of significant connections that the channel participates in).

**S3.5 Network Density Analysis**

To characterize the topology of ripple-associated cortical synchrony, time-resolved functional connectivity networks were constructed from the ripple-locked PLV data. For each participant and 500 ms sliding window, PLV values were averaged across the spindle band (12–16 Hz) and arranged into a symmetric 21×21 adjacency matrix. A fixed absolute threshold set at the top tertile (top 35%) of the PLV distribution pooled across all participants, time windows, and conditions (real and surrogate; threshold = 0.607) was applied uniformly across all time points. Rather than proportional thresholding, which equates network density across time and would suppress genuine fluctuations in synchrony strength, absolute thresholding preserves these differences, with density itself tracked as a primary outcome. This threshold corresponded to the minimum value yielding fully connected graphs (every node in the network can reach every other node through some path of edges), ensuring valid path length and small-world estimation at every time point. Network density was defined as the proportion of surviving edges relative to all possible connections (11): density = $E / (N \times (N-1))$, where E is the number of supra-threshold connections and N = 21. Group-level changes were assessed relative to surrogate controls using cluster-based permutation tests (1000 permutations, two-sided, cluster $\alpha = 0.05$), applied across the full analysis window (−1.0 to 1.0 s).

**S4. Relationship Between Ripple Strength and Cortical Decodability**

To examine whether the strength of hippocampal ripples modulates their detectability in scalp EEG, per-trial LDA decision values (d-values) were correlated with physiological attributes of the corresponding ripple events. This analysis was performed within individual hippocampal contacts to avoid confounding cross-contact variability in baseline ripple properties or recording quality.

For each participant, the mean d-value time course was first smoothed using a 50 ms moving average, and the peak of this smoothed curve was identified within group-level above chance decoding window (−244 to 828 ms relative to ripple peak). D-values were then extracted per trial by averaging across a ±50 ms window centered on the participant-specific peak latency. Three ripple attributes were tested: ripple amplitude, ripple duration, and ripple envelope sum. Spearman rank correlations were computed between the averaged d-values and each attribute separately. To test whether the group-level mean correlation differed significantly from zero while accounting for the non-independence of multiple contacts within the same participant, per-contact Spearman r values were entered into a linear mixed-effects (LME) model with participant as a random intercept (MATLAB fitlme), and the intercept was tested against zero using a t-test on the fixed effect.

Across all 29 contacts, ripple amplitude showed a small but consistent positive correlation with d-values (mean $r = 0.056 \pm 0.075$ SD; median $r = 0.078$), which was reliably above zero at the group level (LME: $\beta = 0.056 \pm 0.015$ SE, $t(28) = 3.805$, $p = 0.0007$). Ripple duration similarly showed a positive association (mean $r = 0.045 \pm 0.075$ SD; median $r = 0.035$; LME: $\beta = 0.045 \pm 0.014$ SE, $t(28) = 3.304$, $p = 0.0026$). The strongest effect was observed for ripple envelope energy (mean $r = 0.062 \pm 0.071$ SD; median $r = 0.063$; LME: $\beta = 0.064 \pm 0.015$ SE, $t(28) = 4.175$, $p = 0.0003$). Together, these results indicate that stronger ripples are more reliably decoded from scalp EEG.

**S5. Cascade Mediation Analysis**

To test whether ripple envelope energy influences cortical synchrony via a sequential pathway through scalp-decodable neural responses and spindle-band power, we implemented a three-link cascade mediation model: ripple envelope energy → scalp decoding strength (d-value at peak discriminability window) → scalp spindle-band power → cortical spindle-band phase synchrony. All four variables were estimated on a per-trial basis and z-scored within contact prior to analysis. Ripple envelope energy and scalp decoding strength are described in S2.2 and S3.3.1, respectively.

Scalp spindle-band power was estimated per trial from the complex Fourier spectra obtained during the ripple-locked time–frequency analysis (S3.1). For each ripple trial, the squared modulus of the Fourier coefficients was averaged across all scalp channels, frequencies within the spindle band (12–16 Hz), and time points spanning 0–0.5 s post-ripple peak, yielding a single scalar power estimate per trial. The 0–0.5 s post-ripple window was motivated by the ripple-locked time–frequency map (Figure 2B), which showed a numerical trend toward elevated t-values within the 12–16 Hz band centered on this period.

Cortical spindle-band phase synchrony was taken from across-scalp PLV (S3.4) during the 500 ms window centered at 300 ms post-ripple (0.05–0.55 s), corresponding to the peak from the group-level cluster analysis. As a control for volume conduction, the cascade was additionally estimated substituting cross-scalp wPLI for cortical PLV as the outcome variable. wPLI was computed across time points within the same 0.05–0.55 s post-ripple window, yielding a single per-trial connectivity value that down-weights phase relationships near zero lag and therefore mitigates effects of volume conduction. Both windows were defined from the observed ripple-locked responses to capture the most physiologically relevant period for each variable.

Each link was estimated as a separate linear mixed-effects model (LME; MATLAB fitlme) with participant and contact as crossed random intercepts: outcome ~ predictor + (1|subId) + (1|macro-contact ID), following the Baron-Kenny causal steps framework adapted for multilevel data (12). The indirect effect was computed as the product of the three path coefficients (a×b×c). Statistical significance of the indirect effect was assessed using a contact-level bootstrap (1,000 iterations), resampling contacts with replacement and refitting all three LME models at each iteration. The 95% confidence interval was taken as the 2.5th–97.5th percentile of the bootstrap distribution; a two-tailed p-value was computed as twice the minimum tail proportion. The total effect (ripple envelope energy → spindle-band PLV, no mediators) and direct effect (controlling for both mediators simultaneously) were also estimated to confirm that ripple envelope energy had no independent influence on cortical synchrony beyond the mediated pathway.

For the primary spindle-band PLV cascade ($n = 12{,}823$ trials), all three links were significant: ripple energy predicted decoding strength ($\beta = 0.067$, SE = 0.009, $t = 7.797$, $p < 0.001$), decoding strength predicted spindle-band power ($\beta = 0.044$, SE = 0.009, $t = 5.072$, $p < 0.001$), and spindle-band power predicted cortical spindle-band PLV ($\beta = 0.095$, SE = 0.009, $t = 11.066$, $p < 0.001$). The indirect effect was significant (a×b×c = 0.0003, 95% CI [0.0001, 0.0005], bootstrap $p = 0.002$). Neither the total effect of ripple energy on cortical PLV nor the direct effect was significant (both ps > 0.663), confirming full sequential mediation.

For the control wPLI cascade, results were consistent: ripple energy predicted decoding strength ($\beta = 0.067$, SE = 0.009, $t = 7.797$, $p < 0.001$), decoding strength predicted spindle-band power ($\beta = 0.044$, SE = 0.009, $t = 5.072$, $p < 0.001$), and spindle-band power predicted cortical spindle-band wPLI ($\beta = 0.040$, SE = 0.009, $t = 4.712$, $p < 0.001$). The indirect effect remained significant (a×b×c = 0.0001, 95% CI [0.0000, 0.0002], bootstrap $p = 0.002$). Neither the total effect ($\beta = -0.009$, $p = 0.279$) nor the direct effect ($\beta = -0.010$, $p = 0.255$) was significant, and the direct path from decoding strength to wPLI controlling for spindle power was also non-significant ($\beta = 0.004$, $p = 0.647$), confirming that the influence of ripple energy on cortical spindle-band wPLI likewise operates through the sequential mediating pathway.

As a confirmatory analysis, the same cascade was estimated using structural equation modeling in R (lavaan (13)), fitting all three paths simultaneously with cluster-robust standard errors (clustered by contact) and percentile bootstrap confidence intervals (1,000 iterations), providing a convergent check on the sequential LME results. For the primary PLV model, the SEM yielded consistent path coefficients (ripple energy → decoding d-value: $\beta = 0.061$, SE = 0.010, $z = 6.36$, $p < 0.001$; decoding d-value → spindle-band power: $\beta = 0.050$, SE = 0.009, $z = 5.32$, $p < 0.001$; spindle-band power → spindle-band PLV: $\beta = 0.100$, SE = 0.010, $z = 10.14$, $p < 0.001$) and a significant indirect effect (a×b×c = 0.0003, 95% CI [0.0002, 0.0005], $z = 3.91$, $p < 0.001$). For the wPLI replication, the SEM similarly confirmed all three links (ripple energy → decoding d-value: $\beta = 0.061$, SE = 0.010, $z = 6.36$, $p < 0.001$; decoding d-value

→ spindle-band power: $\beta = 0.050$, $SE = 0.009$, $z = 5.32$, $p < 0.001$; spindle-band power → spindle-band wPLI: $\beta = 0.041$, $SE = 0.009$, $z = 4.36$, $p < 0.001$) and a significant indirect effect ($a \times b \times c = 0.0001$, 95% CI [0.0001, 0.0002], $z = 3.22$, $p = 0.001$).

#### Figures

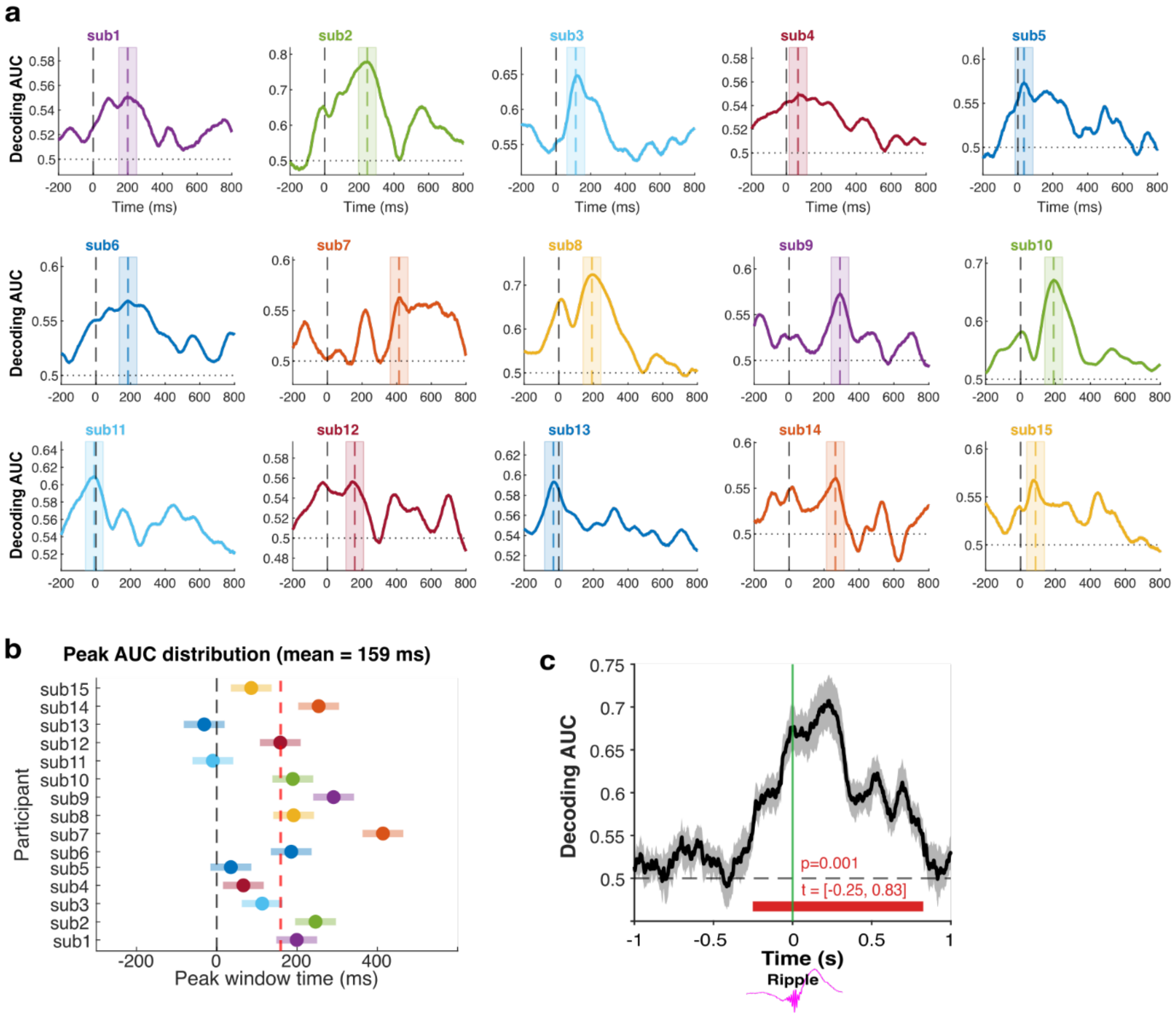


**Fig. S1. Participant-wise decoding AUC time courses and ‘super-trials’ decoding.**

(A) Smoothed classification AUC time courses (50 ms moving average) for each participant (colored lines), plotted relative to hippocampal ripple peak (0 ms, dashed vertical line). Shaded regions indicate the ±50 ms averaging window centered on each participant's peak decoding latency, within which per-trial d-values were averaged for subsequent ripple attribute correlation (S4) and mediation analyses (S5). (B) Participant-specific peak latencies (dots) and corresponding ±50 ms averaging windows (horizontal bars). The red dashed line indicates the group median peak latency. (C) Group-level AUC time course from the ‘super-trial’ analysis (10 trials averaged per ‘super-trial’).

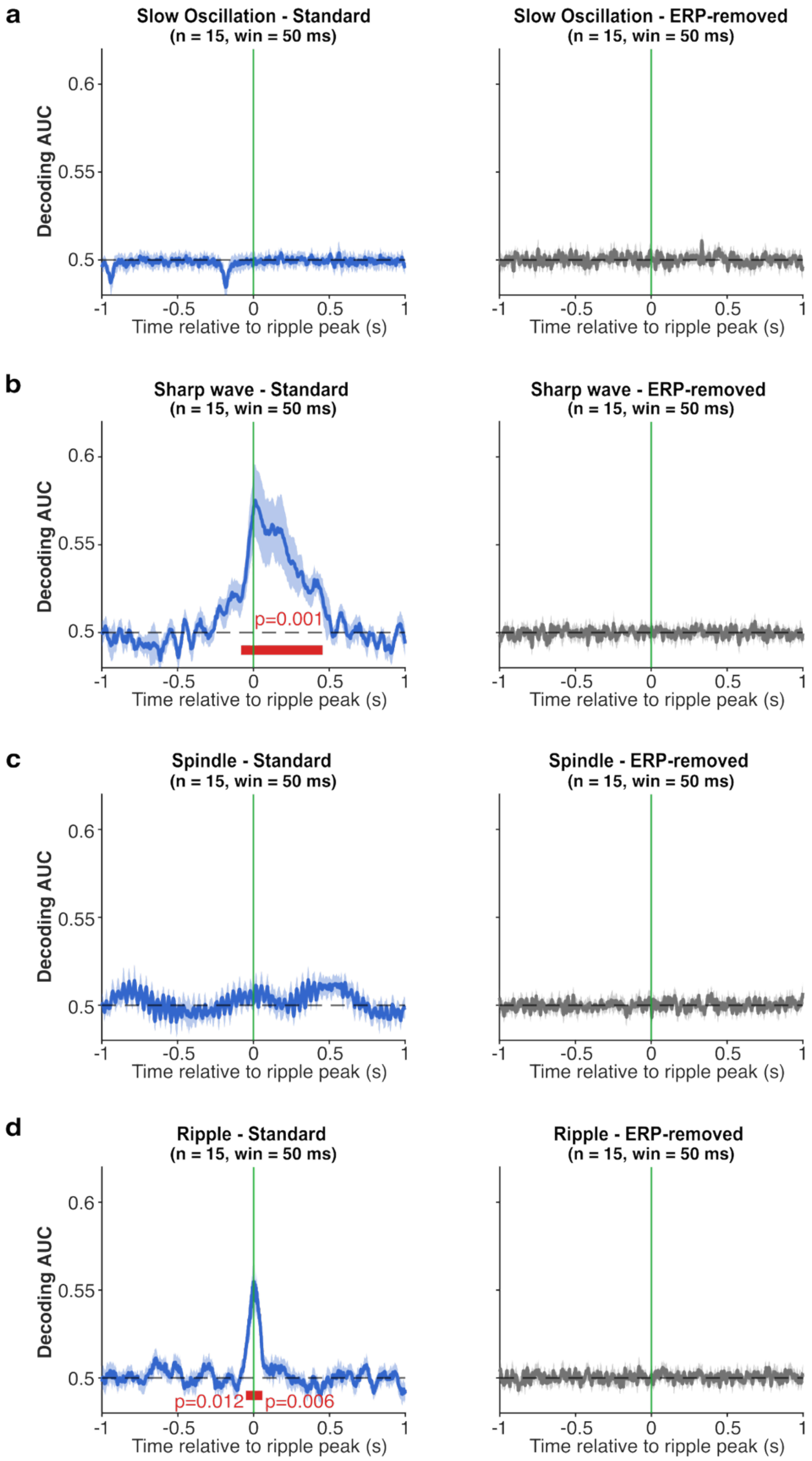


**Fig. S2. Band-specific decoding of ripple-locked scalp EEG signals.**

(A) Slow oscillation band (0.5–1 Hz). Group-level decoding AUC time course (mean ± SEM) for the standard pipeline (left, blue) and ERP-removed control variant (right, grey). (B) Sharp-wave band (2–

9.5 Hz). Significant above-chance decoding was observed from −82 to 456 ms (cluster p = 0.001), with a peak AUC of 0.575 at 12 ms (left). Decoding collapsed to chance following ERP removal (right). (C) Spindle band (12–16 Hz). No significant above-chance decoding was observed in either the standard (left) or ERP-removed (right) variant. (D) Ripple band (80–100 Hz). Significant decoding was confined to two brief peri-event clusters: −52 to 2 ms (p = 0.012) and 6 to 58 ms (p = 0.006; left). Decoding collapsed to chance following ERP removal (right). For all panels, the vertical green line marks ripple peak (t = 0); the dashed horizontal line indicates chance level (AUC = 0.5); red horizontal bars indicate significant above-chance clusters.

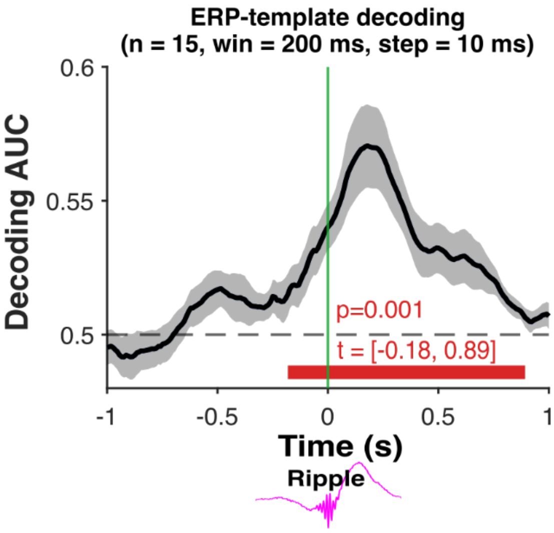


**Fig. S3. ERP-Template Decoding of ripple-locked scalp EEG signals.**
Traces show group mean ± SEM; dashed horizontal line indicates chance level (AUC = 0.5); vertical line marks ripple peak (t = 0).

#### Tables

**Table S1.** Participant demographic and recording information. Demographic and hippocampal contact information reported for each participant (Sub 1-15), including sex (male/female), age at time of recording (years), total number of channels included in the analyses.

| Sub | Sex | Age | Handedness | # Scalp channels | # Hippocampal macro-contacts | |
|---|---|---|---|---|---|---|
| | | | | | L | R |
| 1 | M | 21 | R | 21 | 2 | 0 |
| 2 | M | 20 | R | 21 | 0 | 1 |
| 3 | M | 30 | R | 21 | 0 | 2 |
| 4 | F | 20 | R | 21 | 0 | 2 |
| 5 | F | 47 | R | 21 | 2 | 0 |
| 6 | F | 58 | L | 21 | 1 | 1 |
| 7 | F | 23 | R | 21 | 2 | 0 |
| 8 | M | 39 | R | 21 | 0 | 2 |
| 9 | M | 19 | L | 19 | 0 | 2 |
| 10 | F | 35 | R | 21 | 0 | 2 |
| 11 | F | 24 | R | 21 | 2 | 0 |
| 12 | F | 35 | R | 21 | 2 | 0 |
| 13 | M | 27 | R | 21 | 2 | 0 |
| 14 | M | 21 | R | 21 | 0 | 2 |
| 15 | F | 25 | R | 21 | 2 | 0 |

**Table S2.** Hippocampal macro-contact locations and artifact-free sleep duration per participant. MNI coordinates correspond to the most medial hippocampal macro-contact on each electrode shaft. Sleep duration reflects artifact-free time within N2 and N3, following automated artifact rejection described in S2.1.

| | | MNI coordinate | | | Sleep duration (min) | |
|---|---|---|---|---|---|---|
| **Sub** | **Contact** | **x** | **y** | **z** | **N2** | **N3** |
| 1 | LTA1 | -22 | -11 | -21 | 186.0 | 83.8 |
| | LTB1 | -28 | -23 | -14 | 197.4 | 94.3 |
| 2 | RTA1 | 25 | -18 | -17 | 98.9 | 50.1 |
| 3 | RTB1 | 32 | -12 | -23 | 137.2 | 33.0 |
| | RTC1 | 32 | -21 | -13 | 161.3 | 49.5 |
| 4 | RTA1 | 34 | -10 | -20 | 222.5 | 160.8 |
| | RTB1 | 32 | -23 | -9 | 205.4 | 128.8 |
| 5 | LTA1 | -24 | -14 | -23 | 214.5 | 0.0 |
| | LTB1 | -33 | -26 | -18 | 162.4 | 0.0 |
| 6 | LTA1 | -32 | -20 | -16 | 241.8 | 71.6 |
| | RTC1 | 36 | -19 | -14 | 212.2 | 56.6 |
| 7 | LTB1 | -27 | -17 | -25 | 135.1 | 84.4 |
| | LTD1 | -29 | -26 | -12 | 135.7 | 84.7 |
| 8 | RTC1 | 27 | -20 | -25 | 103.8 | 74.0 |
| | RTE1 | 27 | -29 | -20 | 106.7 | 75.7 |
| 9 | RTC1 | 28 | -22 | -14 | 124.2 | 91.6 |
| | RTB1 | 28 | -13 | -22 | 117.7 | 86.0 |
| 10 | RTC1 | 33 | -26 | -14 | 183.1 | 38.1 |
| | RTB1 | 34 | -15 | -23 | 142.9 | 25.6 |
| 11 | LTC1 | -29 | -12 | -16 | 98.6 | 36.2 |
| | LTE1 | -34 | -22 | -10 | 98.4 | 35.5 |
| 12 | LTD1 | -29 | -23 | -14 | 152.2 | 57.7 |
| | LTB1 | -29 | -13 | -22 | 123.7 | 44.5 |
| 13 | RTC1 | 31 | -22 | -13 | 161.1 | 76.4 |
| | RTB1 | 32 | -12 | -22 | 153.9 | 71.7 |
| 14 | RTA1 | 28 | -15 | -16 | 158.7 | 26.0 |
| | RTB1 | 36 | -22 | -11 | 176.0 | 29.3 |
| 15 | LTC1 | -30 | -25 | -19 | 118.6 | 43.8 |
| | LTD1 | -27 | -39 | -8 | 130.2 | 46.2 |

**Table S3.** Hippocampal sharp-wave ripple attributes during NREM sleep per recording contact. N = total NREM ripple count (N2 + N3); Density = total NREM ripples per artifact-free minute; Duration = mean ripple duration (ms); Amplitude = mean peak amplitude (μV); Frequency = mean instantaneous ripple-band frequency (Hz); Envelope sum = mean summed ripple envelope energy, reflecting overall ripple strength. Summary statistics (mean ± SD) are computed across participants (n = 15).

| | | NREM ripple attributes | | | | | |
|---|---|---|---|---|---|---|---|
| **Sub** | **Contact** | **N** | **Density (min)** | **Duration (ms)** | **Amplitude (μV)** | **Frequency (Hz)** | **Envelope sum** |
| 1 | LTA1 | 694 | 2.57 | 71.3 | 7.55 | 87.8 | 148.4 |
| | LTB1 | 462 | 1.58 | 57.8 | 11.30 | 88.7 | 200.4 |
| 2 | RTA1 | 95 | 0.64 | 60.6 | 8.49 | 97.3 | 132.4 |
| 3 | RTB1 | 100 | 0.59 | 68.7 | 12.54 | 88.4 | 235.2 |
| | RTC1 | 215 | 1.02 | 71.6 | 6.12 | 92.1 | 119.9 |
| 4 | RTA1 | 2403 | 6.27 | 63.3 | 7.44 | 89.2 | 141.4 |
| | RTB1 | 858 | 2.57 | 61.2 | 4.42 | 88.9 | 82.9 |
| 5 | LTA1 | 231 | 1.08 | 54.6 | 6.29 | 89.2 | 105.4 |
| | LTB1 | 228 | 1.40 | 53.6 | 5.26 | 90.9 | 86.5 |
| 6 | LTA1 | 957 | 3.05 | 67.0 | 4.59 | 93.3 | 88.7 |
| | RTC1 | 1174 | 4.37 | 74.2 | 2.18 | 89.3 | 45.6 |
| 7 | LTB1 | 260 | 1.18 | 65.3 | 2.19 | 95.7 | 41.0 |
| | LTD1 | 175 | 0.79 | 59.1 | 1.82 | 94.1 | 31.7 |
| 8 | RTC1 | 388 | 2.18 | 65.4 | 7.35 | 89.5 | 129.4 |
| | RTE1 | 147 | 0.81 | 58.4 | 3.49 | 90.4 | 58.0 |
| 9 | RTC1 | 394 | 1.83 | 56.3 | 5.69 | 91.3 | 96.0 |
| | RTB1 | 273 | 1.34 | 60.5 | 8.03 | 89.5 | 141.1 |
| 10 | RTC1 | 995 | 4.50 | 61.3 | 12.28 | 91.8 | 221.5 |
| | RTB1 | 1524 | 9.04 | 71.6 | 11.28 | 93.3 | 217.4 |
| 11 | LTC1 | 584 | 4.33 | 59.3 | 6.37 | 89.4 | 114.1 |
| | LTE1 | 203 | 1.52 | 57.5 | 3.90 | 90.0 | 68.4 |
| 12 | LTD1 | 394 | 1.88 | 61.0 | 3.21 | 89.4 | 57.2 |
| | LTB1 | 330 | 1.96 | 70.2 | 6.51 | 87.8 | 126.5 |
| 13 | RTC1 | 1006 | 4.24 | 59.6 | 4.94 | 86.6 | 90.5 |
| | RTB1 | 598 | 2.65 | 57.8 | 3.50 | 86.8 | 61.8 |
| 14 | RTA1 | 325 | 1.76 | 64.1 | 3.17 | 91.4 | 59.3 |
| | RTB1 | 127 | 0.62 | 50.1 | 2.09 | 93.9 | 32.3 |
| 15 | LTC1 | 354 | 2.18 | 66.3 | 4.54 | 90.9 | 84.2 |
| | LTD1 | 385 | 2.18 | 76.3 | 3.32 | 90.5 | 70.1 |
| Mean ± SD | | 532 ± 455 | 2.36 ± 1.69 | 63.5 ± 5.3 | 5.81 ± 2.81 | 90.9 ± 2.6 | 107.4 ± 52.6 |
